# NSD2 overexpression drives clustered chromatin and transcriptional changes in a subset of insulated domains

**DOI:** 10.1101/587931

**Authors:** Priscillia Lhoumaud, Sana Badri, Javier Rodriguez Hernaez, Theodore Sakellaropoulos, Gunjan Sethia, Andreas Kloetgen, MacIntosh Cornwell, Sourya Bhattacharyya, Ferhat Ay, Richard Bonneau, Aristotelis Tsirigos, Jane A. Skok

## Abstract

CTCF and cohesin play a key role in organizing chromatin into TAD structures. Disruption of a single CTCF binding site is sufficient to change chromosomal interactions leading to alterations in chromatin modifications and gene regulation. However, the extent to which alterations in chromatin modifications can disrupt 3D chromosome organization leading to transcriptional changes is unknown. In multiple myeloma a 4;14 translocation induces overexpression of the histone methyltransferase, NSD2 resulting in expansion of H3K36me2 and shrinkage of antagonistic H3K27me3 domains. Using isogenic cell lines producing high and low levels of NSD2, we find oncogene activation is linked to alterations in H3K27ac and CTCF within H3K36me2 enriched chromatin. A linear regression model reveals that changes in both CTCF and/or H3K27ac significantly increase the probability that a gene sharing the same insulated domain will be differentially expressed. These results identify a bidirectional relationship between 2D chromatin and 3D genome organization in gene regulation.

## Introduction

The 3D organization of chromosomes enables cells to balance the biophysical constraints of the crowded nucleus with the functional dynamics of gene regulation. Chromosomes are divided into large domains that physically separate into two nuclear compartments (A and B) of active and inactive chromatin, respectively ^1^. Loci within each compartment interact more frequently with other loci from the same compartment irrespective of whether these regions are proximal on the linear chromosome. Compartments can be further subdivided into 1MB sized highly self-interacting ‘topologically associated domains’ (TADs), which are separated from each other by insulating boundaries enriched for the architectural proteins, CTCF and cohesin ^2–4^. Since TADs are largely conserved between cell types and across different species they are considered the basic organizational unit of eukaryotic genomes. They play an important role in gene expression by limiting the influence of enhancers to genes located within the same TAD ^5–7^.

CTCF and cohesin are major contributors in shaping architecture within TADs and depletion of either factor leads to loss of TAD structure ^8–10^. Depletion of cohesin also leads to strengthening of compartments, suggesting that compartments and TADs are formed by two different mechanisms and that TAD structures antagonize compartmentalization by mixing regions of disparate chromatin status ^9,10^. The best-accepted model to explain TAD formation and maintenance, involves a loop-extrusion mechanism whereby cohesin rings create loops by actively extruding DNA until the complex finds two CTCF binding sites in convergent orientation ^11^. In this way cohesin forms chromatin loops as a result of its ability to hold together two double-strand DNA helices via its ring structure ^11–15^. Indeed, genome-wide analyses revealed that loops are preferentially formed between convergently orientated CTCF binding sites ^16,17^ and divergent sites delineate boundary regions ^18^. These sites are thought to serve as a block to the movement of cohesin on chromatin.

A number of labs have demonstrated that disruption of a single CTCF binding site is sufficient to alter chromosomal interactions leading to the spreading of active chromatin and altered gene regulation ^14,19–21^. However, it is not known if the reverse is true and whether alterations in chromatin modifications themselves can impact chromosome organization at the level of A and B compartments, TAD structure, CTCF binding and enhancer-promoter contacts in a manner that corresponds to changes in gene regulation. An alteration in the balance of the antagonistic marks H3K36me2 and H3K27me3 is a hallmark of many different cancers. Translocation between the immunoglobulin heavy chain locus, *IGH* on chromosome 14 with the *NSD2* locus (also known as MMSET or WHSC1) on chromosome 4, leads to NSD2 overexpression in 15-20% of multiple myeloma (MM) patients that have a poor survival rate and do not respond well to cytotoxic chemotherapy ^22–25^. NSD2 is a histone methyl transferase (HMT) that is responsible for deposition of the H3K36 mono- and di-methyl mark. In a wild-type setting, H3K36me2 accumulates on active gene bodies and acts as a signature of transcriptional activity. However, when NSD2 is overexpressed as a result of the 4;14 translocation, H3K36me2 spreads outside of active gene bodies into intergenic regions. Expansion of H3K36me2 domains results in contraction of H3K27me3 domains, altering gene expression programs in the absence of driver mutations in a manner that is poorly understood ^26^.

Of note, similar changes in chromatin are detected in other cancers such as B and T acute lymphoblastic leukemia (B- and T-ALL) and a number of advanced stage solid tumors, including prostate, colon and skin cancers ^27,28^. Increased H3K36me2 in some cases can result from an E1099K mutation in NSD2 that affects the catalytic domain of this enzyme ^29^. In two pediatric brain cancers, diffuse intrinsic pontine glioblastoma (DIPG) and supratentorial glioblastoma multiforme (GBMs), a mutation in H3.3 in which the lysine at position 27 is mutated to a methionine (H3K27M) results in a similar H3K36me2 versus H3K27me3 imbalance by impacting the action of EZH2 ^30–32^. DIPG and GBM are the most aggressive primary malignant brain tumors in adults and children and the median survival of this group of patients is approximately 1 year. Given the poor prognosis of patients suffering from these different cancers, it is important to gain a better understanding of the mechanisms underlying changes in gene expression in diseases with an H3K36me2 versus H3K27me3 imbalance.

In multiple myeloma, alterations in gene expression have been shown to be dependent on the histone methyl-transferase activity of NSD2 ^33^. Although the impact of NSD2 overexpression on chromatin modifications has been well documented, there has been no in-depth analysis into the mechanisms underlying the changes in gene expression that occur downstream of the expansion and reduction of active H3K36me2 and repressive H3K27me3 domains. Using isogenically matched multiple myeloma patient derived cell lines that differ only in the levels of NSD2 they express, we demonstrate that spreading of H3K36me2 from active gene bodies into intergenic regions is accompanied by changes in H3K27ac (a feature of regulatory elements) as well as an alteration in the genome wide binding profile of CTCF. Both changes are linked to significant alterations in gene expression and oncogene activation. Expansion of H3K36me2 domains also drives compartment switching and alterations in intra-TAD interactions, while altered boundary insulation scores overlap differential CTCF and Rad21 binding. Importantly, analysis of Hi-C and CTCF HiChIP data demonstrates that changes in CTCF and H3K27ac cluster together with alterations in transcriptional output within a subset of TADs and CTCF loops in NSD2 high expressing cells. A linear regression model reveals that changes in both CTCF and/or H3K27ac significantly increases the probability that a gene sharing the same insulated domain will be differentially expressed. These results demonstrate a bidirectional relationship between 2D and 3D chromatin organization in gene regulation and demonstrate that cells can co-opt altered chromatin domains to drive oncogenic transcriptional programs that are regulated within insulated boundaries.

## Results

### NSD2 overexpression leads to alterations in H3K27ac

NSD2 overexpression leads to spreading of H3K36me2 from active gene bodies into intergenic regions leading to a more open chromatin conformation ^24^. Alterations in gene expression have previously been shown to be dependent on the histone methyl-transferase activity of NSD2, highlighting the importance of deposition and expansion of H3K36me2 domains in the activation of oncogenic transcriptional pathways ^33^. However, it is not known how intergenic spreading of H3K36me2 alters gene regulation and whether 3D organization of the genome plays a role.

In order to address this question, we used two isogenic cell lines generated from a patient derived KMS11 t(4;14) multiple myeloma cell line: NTKO (non-translocated knockout) and TKO (translocated knockout) cells, which have the translocated allele or the endogenous *NSD2* allele, respectively inactivated (Fig. 1a) ^25^. Importantly, NTKO and TKO cells differ solely in the level of NSD2 they express and henceforth are referred to as NSD2 High and Low cells. The paired cell lines provide us with an opportunity to analyze the impact of NSD2 overexpression independent of other confounding genetic or epigenetic alterations. In this respect they are more useful than patient-based samples which do not have appropriate controls. Using RNA-seq we observed that NSD2 High expression leads to the deregulation of many genes (1650 up and 303 downregulated genes) (Fig. 1b, **Supplementary Table 1**) and principal component analysis (PCA) revealed that NSD2 High and Low replicates separated into distinct clusters (**Supplementary Fig. 1a**), a finding that is consistent with previous studies ^24,26^. Genes associated with multiple myeloma and KRAS pathways were enriched as shown by Gene Set Enrichment Analysis (GSEA, **Supplementary Fig. 1b**)

**Fig. 1.**
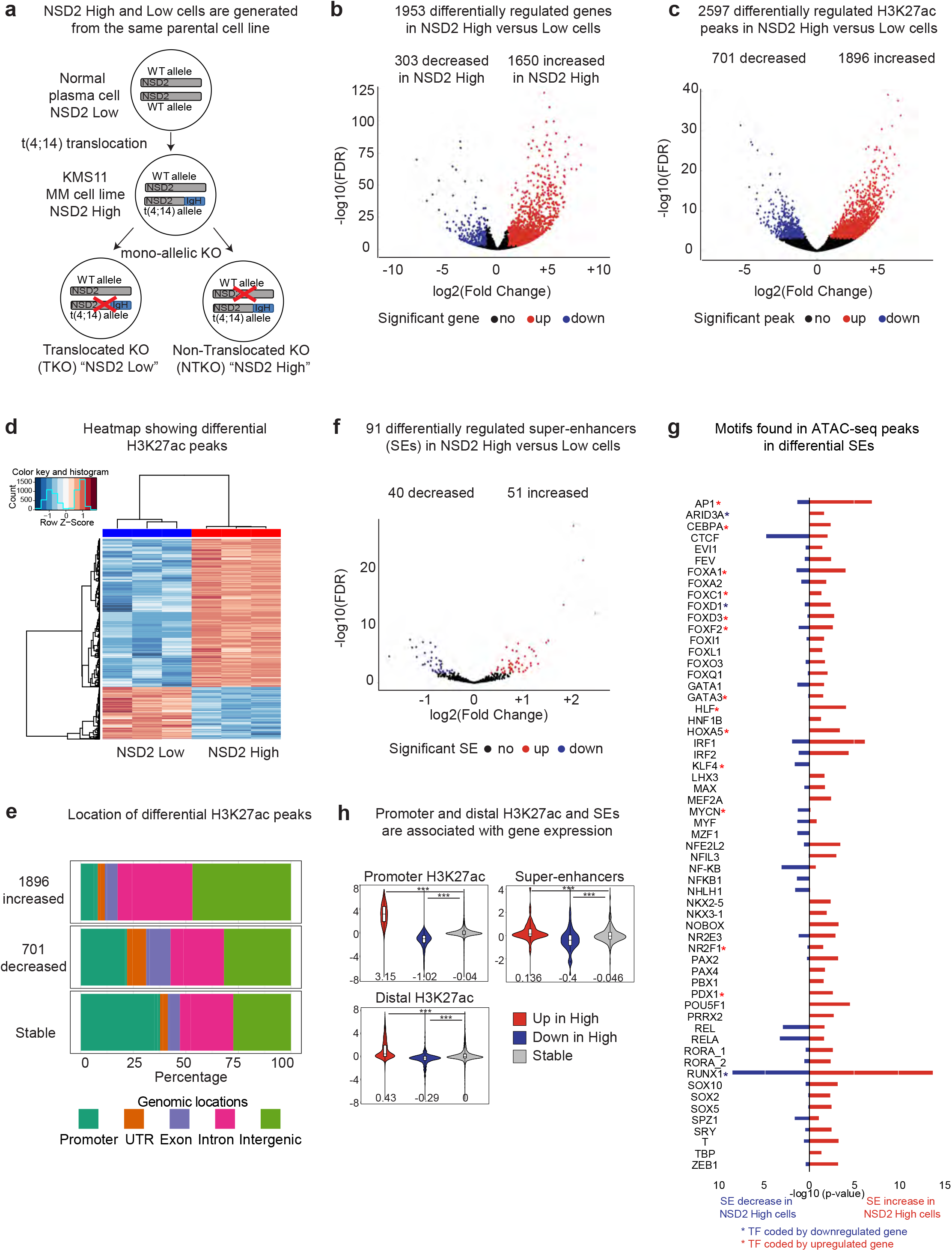
NSD2 overexpression leads to alterations in H3K27ac enrichment linked to changes in gene expression. **a**, NSD2 Low and High isogenic cell lines generated from a patient derived KMS11 t(4;14) multiple myeloma cell line: NTKO (non-translocated knockout) and TKO (translocated knockout) cells, which have the translocated allele or the endogenous *NSD2* allele, respectively inactivated. **b**, Volcano plot showing significant NSD2-mediated changes in gene expression (FDR <0.01). Upregulated genes: 1650 (red, log2 fold change >1); downregulated genes: 303 (blue, log2 fold change <-1). **c**, Volcano plot showing significant NSD2-mediated changes in H3K27ac (FDR <0.01). Decreased peaks: 701 (blue, log2 fold change <-1); increased peaks: 1896 (red, log2 fold change >1). **d**, Heatmap showing differential H3K27ac peaks identified using DiffBind analysis of ChIP-seq triplicates in NSD2 High versus Low cells. Increase and decrease in gene expression is shown in red and blue, respectively. **e**, Genomic locations of the differential H3K27ac peaks. **f**, Volcano plot showing significant NSD2-mediated changes in super-enhancers (FDR <0.01). Super-enhancers in NSD2 High versus Low cells were called based on H3K27ac levels using ROSE. Increased super-enhancers: 51 (red) and decreased super-enhancers: 40 (blue). **g**, Transcription factor motifs identified in 119 ATAC-seq peaks of increased (51) and decreased (40) super-enhancers using TRAP. Motifs found in increased (red) and decreased (blue) super-enhancers (-log10 p-value). Red and blue stars indicate that the gene coding for the TF is respectively up-or downregulated in NSD2 High cells. **h**, Gene expression changes are associated with H3K27ac changes at promoters (top panel, -/+ 3kb around TSS), distal H3K27ac and super-enhancers (middle and bottom panels, respectively, 3-250kb up and downstream from TSSs associated with genes using GREAT). H3K27ac up, down and stable red, blue and grey in NSD2 High versus Low cells.

Since the vast majority of regulatory elements are located within intergenic regions, we asked whether spreading of H3K36me2 into these locations could impact their activation status. To test this idea, we analyzed H3K27ac, a feature of both enhancers and promoters. H3K27ac ChIP-seq revealed that a total of 2597 peaks were significantly affected by NSD2 overexpression and PCA revealed that NSD2 High and Low replicates clustered separately (**Supplementary Fig. 2a**). Specifically, we identified 1896 increased and 701 decreased peaks in NSD2 High versus low cells (Fig. 1c, d, **Supplementary Table 2**). Increased and decreased peaks were predominantly located in intergenic and intronic regions suggesting that enhancer activity could be affected (Fig. 1e). This pattern is much more pronounced for increased peaks.

Previous studies have reported changes in the activity of super-enhancers in multiple myeloma ^34,35^. The term super enhancer is used to describe clusters of enhancers with high levels of activating histone marks including H3K27ac ^36^. Super-enhancers are occupied by lineage specific master regulator transcription factors (TFs) known to control cell identity ^36,37^. Furthermore, super-enhancers are generated at oncogenes and other loci that are important in tumorigenesis ^36,37^. To separate super-enhancers from typical enhancers we used ‘ROSE’ (rank ordering of super-enhancers ^36^). With this approach, we identified 260 and 278 super-enhancers in NSD2 low and high cells, respectively ^35,37^ (**Supplementary Fig. 2b**). Differential super-enhancers were identified from the merged group based on alterations in H3K27ac enrichment using an FDR cutoff of 0.1. With this strategy, we identified about a third of super-enhancers (91) with significantly altered H3K27ac signal in NSD2 High cells, 51 with increased and 40 with decreased signal (Fig. 1f).

### Changes in H3K27ac are associated with changes in gene regulation

To determine which transcription factors (TFs) could bind gained and lost super-enhancers in NSD2 High versus Low cells, we analyzed motifs using ATAC-seq. We identified 119 ATAC-seq peaks in the 51 gained super-enhancers and 102 ATAC-seq peaks in the 40 super-enhancers that were lost (Fig. 1g). TRAP (transcription factor affinity prediction) was used to identify which motifs were present in the ATAC-seq peaks ^38^. Some of the motifs were shared between gained and lost super-enhancers (CTCF, RUNX1 etc), while others were identified in only one category. Furthermore, we found that the expression of some of the genes that encode these TFs goes up or down as denoted respectively by a red or blue asterisk in Fig. 1f, Most of the motifs in super-enhancers were also found in individual gained and lost H3K27ac peaks, with the exception of the CTCF motif (**Supplementary Fig. 2c**). This is consistent with recent studies showing that super-enhancers are enriched with CTCF more frequently than typical enhancers ^39,40^.

To examine the connections between alterations in H3K27ac and gene regulation we separated H3K27ac peaks into those located at promoters, non-promoter sites (intragenic and intergenic combined) and super-enhancers. GREAT (Genomic Regions Enrichment of Annotations Tool; ^41^) was used to associate peaks at distal sites and super-enhancers with gene expression changes, using an arbitrary cutoff of 250kb, based on findings from a previous study showing that the mean distance from enhancer to promoter is around 196kb ^42^. H3K27ac peaks at all three sites were significantly correlated with gene expression changes (Fig. 1h), with the strongest effect seen at promoters, possibly because this is the most accurate prediction. Together these data indicate that overexpression of NSD2 leads to significant changes in H3K27ac linked to deregulation of gene expression.

### Changes in CTCF binding profile are linked to alterations in gene expression

Nuclear architecture is a powerful regulator of gene expression. In particular, precise transcriptional control is exerted by restricting the influence of enhancers to target genes within TADs whose boundaries are enriched for the insulating protein, CTCF. Given that marks associated with regulatory elements are significantly altered in a manner that corresponds to changes in gene expression in NSD2 overexpressing cells, we next asked whether alterations in histone modifications could lead to changes in chromosome organization. For this analysis, we started with a CTCF ChIP-seq and identified a surprising difference in CTCF binding: 2197 and 295 CTCF peaks were increased and decreased, respectively in the NSD2 High cells (Fig. 2a and 2b, **Supplementary Table 3**) that separated into distinct clusters by PCA analysis in NSD2 High and Low replicates (**Supplementary Fig. 2d**). Increased CTCF peaks were predominantly located in intergenic and intragenic regions, as compared with decreased and stable CTCF peaks (Fig. 2c). Moreover, CTCF peaks that were enriched or depleted at promoters and distal sites were significantly correlated with gene expression changes (Fig. 2d), with the strongest effect seen at promoters. This finding is consistent with previous studies showing that loss of CTCF at promoters leads to downregulation of genes ^8^.

**Fig. 2.**
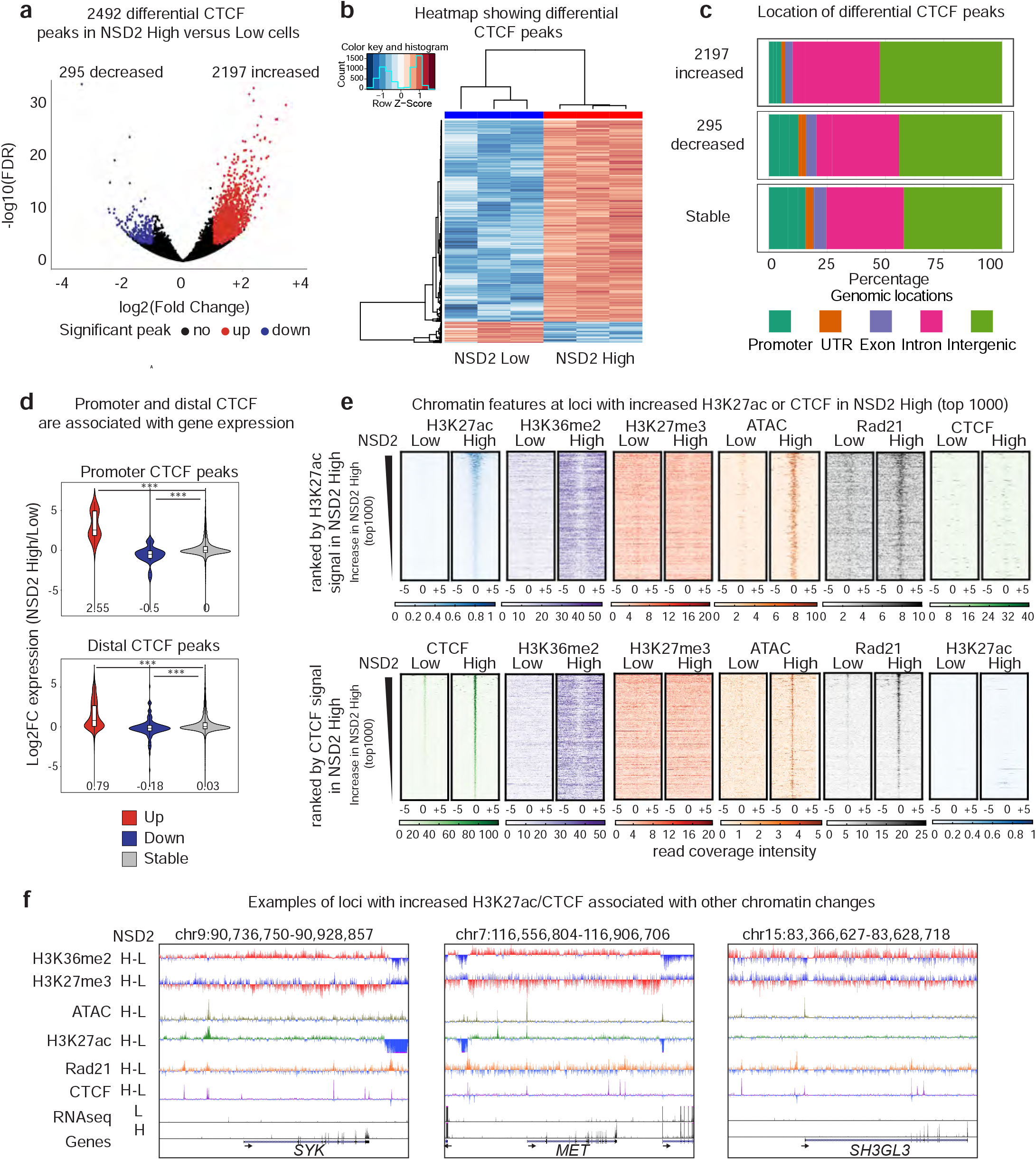
New CTCF and H3K27ac peaks are located within expanded H3K36me2 domains. **a**, Volcano plot showing significant NSD2-mediated changes in CTCF binding (FDR <0.01). Increasing peaks: 1650 (red, log2 fold change >1); decreasing peaks: (blue, log2 fold change <-1). **b**, Heatmap showing differential CTCF peaks identified using DiffBind analysis of ChIP-seq triplicates in NSD2 High versus Low cells. Increases and decreases in gene expression are shown in red and blue, respectively. **c**, Genomic locations of differential CTCF peaks. **d**, Gene expression changes are associated with CTCF changes at promoters (top panel, −/+ 3kb around TSS) and distal CTCF (bottom panel, 3-250kb up and downstream from TSSs associated with genes using GREAT). CTCF up, down and stable red, blue and grey in NSD2 High versus Low cells. **e**, Heatmaps of H3K27ac, H3K36me2, H3K27me3, ATAC-seq, Rad21 and CTCF signal at the top 1000 increased H3K27ac (top panels) and CTCF (bottom panels) peaks in NSD2 High cells. Peaks are ranked by H3K27ac (top panels) and CTCF (bottom panels) signal in NSD2 High cells. **f**, UCSC genome browser screenshots of chromatin features at regions surrounding three oncogenes (*SYK*, left panel; *MET*, middle panel; and *SH3GL3*, right panel).

### New CTCF and H3K27ac peaks are located within expanded H3K36me2 domains

To better understand the links between NSD2 overexpression and the altered genomic profiles of H3K27ac and CTCF, we analyzed the chromatin landscape surrounding the 1000 most enriched new H3K27ac and CTCF peaks in the NSD2 High cells. Importantly, new H3K27ac and CTCF peaks were found to be located in newly enriched H3K36me2 regions while H3K27me3 remained unchanged at these locations (Fig. 2e). Thus, increases in H3K27ac peaks were found to be independent of H3K27me3 changes. New H3K27ac and CTCF peaks were also associated with a gain in overlapping ATAC-seq peaks. This data suggests that spreading of H3K36me2 provides a more accessible chromatin landscape favorable for CTCF and H3K27ac enrichment. As shown in Fig. 2f, these changes are linked to the activation of oncogenes involved in multiple myeloma including *SYK* ^43–47^*, MET* ^48–50^ and *SH3GL3* ^51^ in NSD2 overexpressing cells (Fig. 2f). Of note, few changes were observed for the 1000 most stable and decreased H3K27ac and CTCF peaks, but a decrease in both H3K27ac and CTCF was linked to a slight increase in H3K27me3 (**Supplementary Fig. 3**).

The vast majority of chromosomal loops involve cohesin. These can be separated into two main categories: loops that are CTCF dependent and CTCF independent, cell type specific, dynamic loops that form between enhancers and promoters and involve cell type specific TFs ^52^. To investigate whether new H3K27ac and CTCF peaks had the potential to be involved in looping in the NSD2 High cells we performed ChIP-seq for Rad21 (a component of the cohesin complex). Interestingly, new Rad21 peaks were associated with both newly enriched H3K27ac and CTCF sites in the NSD2 overexpressing cells. However, gained CTCF and H3K27ac peaks did not overlap with each other (Fig. 2e), suggesting that loops involving new regulatory elements associated with enriched H3K27ac were generated in a CTCF independent manner.

### NSD2 overexpression drives A/B compartment switching

Given the finding that there are significant changes in CTCF or H3K27ac peaks overlapping Rad21 alterations in NSD2 High cells, we next asked whether 3D organization was altered. To determine this, we performed Hi-C in duplicate using the Arima Kit and processed the data using Hi-C bench ^53^. The number of valid read pairs was consistent between replicates (~120 million to ~180 million (**Supplementary Fig. 4a**)) and the PCA showed that NSD2 High versus Low replicates separated as expected (**Supplementary Fig. 4b**). A/B compartment status was analyzed in merged NSD2 Low and High replicates using the Eigen vector (principal component 1, PC1, see Supplementary Information for details) ^1^. We observed significant compartment weakening in NSD2 High cells, as exemplified by the changes visualized on chromosome 7 in Fig. 3a. Overall, the number of A and B compartments was lower in NSD2 High versus Low cells and we detected 324 regions that had switched from A to B and 491 regions that switched from B to A (Fig. 3b-d). Although there were more regions switching from B to A, these regions were smaller and the portion of the genome was comparable to the regions switching from A to B (**Supplementary Fig. 4c-e**). Compartment switching from B to A overlapped with expanded H3K36me2 domains and a reduction in H3K27me3, while the reverse was true for switching from compartment A to B (Fig. 3d, e). We speculate that an overall increase in CTCF peaks in the genome in NSD2 High cells could contribute to weakening of compartment structure, consistent with previous findings showing that loss of cohesin and TAD structure strengthens compartmentalization ^9,10^.

**Fig. 3.**
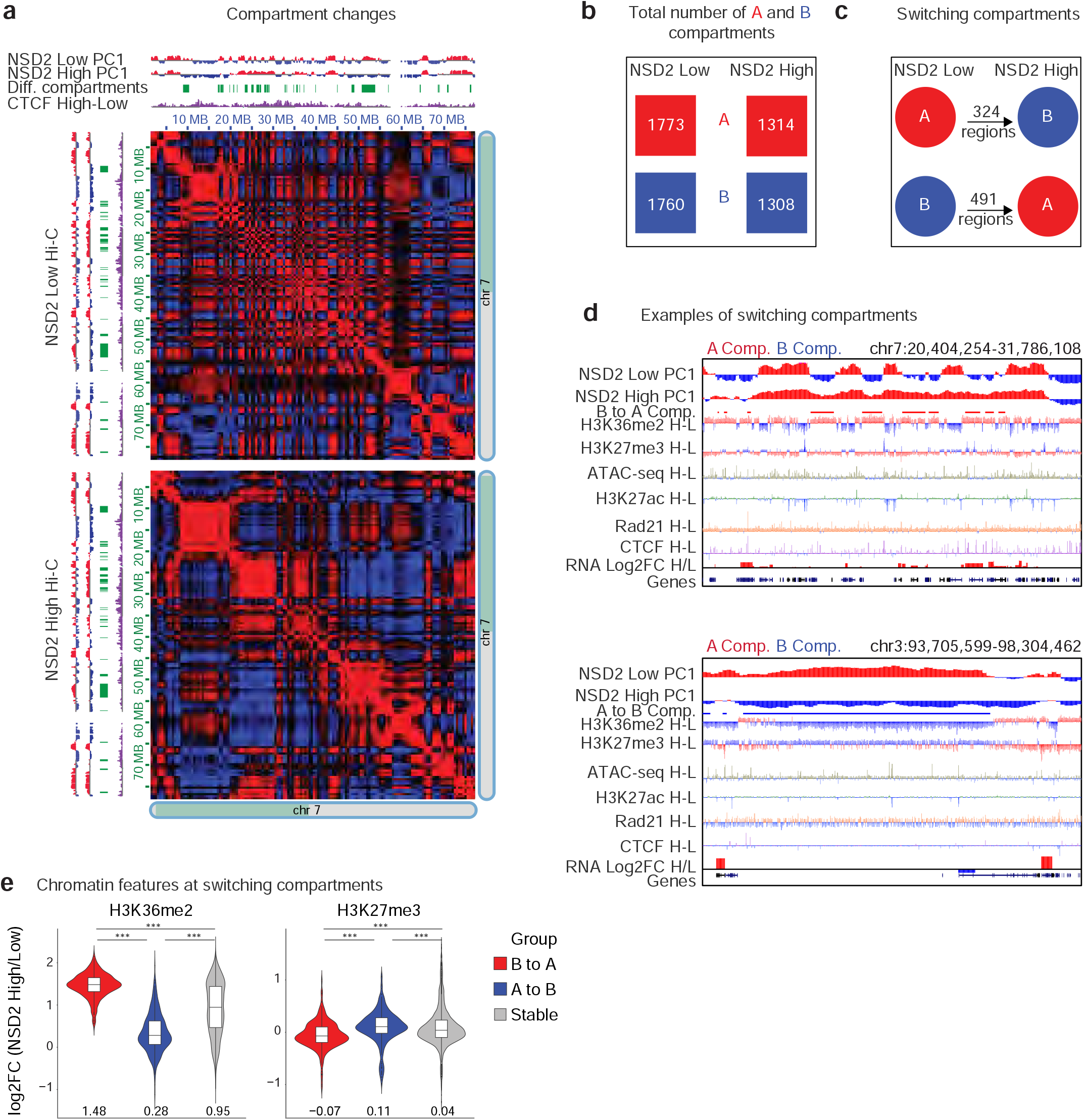
NSD2 overexpression drives A/B compartment switching. **a**, Compartment weakening for half of chromosome 7 is shown in NSD2 High versus Low cells. Top and right: Eigen vector (PC1) for compartments A and B in red and blue, respectively. Switching regions are shown in green and in purple the subtraction of CTCF signal (NSD2 High - Low). Heatmaps represent the Pearson correlation of interactions in NSD2 Low (top) and High (bottom) cells. Positive and negative Pearson correlation between two loci are represented in red and blue, respectively. **b**, Total number of A (red) and B (blue) compartments in NSD2 High versus Low cells. **c**, Number of regions that switch compartments from B to A in (491 regions) or A to B (324 regions) in NSD2 High versus Low cells. **d**, IGV screenshots show examples of regions that switch from A to B (top panel) and B to A (bottom panel) in NSD2 High versus Low cells. Eigen vectors (PC1), differential compartment switching, subtraction tracks of H3K36me2 and H3K27ac and expression for NSD2 High and Low cells are shown. Regions that significantly switch from B to A and from A to B are indicated in red (top panel, B to A) or in blue (bottom panel, A to B), respectively. **e**, Changes in H3K36me2 and H3K27me3 levels within regions that switch compartment from B to A (red), A to B (blue) or are stable (grey) in NSD2 High cells. Log2 fold changes (NSD2 High versus Low cells) for H3K36me2 and H3K27me3 are shown. The median is indicated under the violins.

### Changes in TAD boundaries and intra-TAD interactions

Both Hi-C replicates from each condition harbored similar number of TADs with similar sizes (**Supplementary Fig. 4f, g**) and they were merged for TAD analysis to determine the impact of NSD2 overexpression on 3D genome organization. Using Hi-C bench, we assigned a mean boundary score to each replicate using a resolution of 40kb ^53^ to determine if there were changes in TAD boundaries (see Supplementary Information for details). NSD2 overexpression was associated with a significant (FDR < 0.05) increase in insulation at TAD boundaries (red, 61) and much fewer boundaries that were weakened (blue, 5) (Fig. 4a). These changes were synonymous with changes in CTCF and Rad21 binding (Fig. 4b). In addition, we found that NSD2 High cells were predominantly associated with significant gains (red, 229) and fewer decreases in intra-TAD interactions (blue, 30) (Fig. 4c). Increased intra-TAD interactions were linked to expanded H3K36me2 domains and a reduction in H3K27me3, while decreases in intra-TAD interactions were linked to a reduction in H3K36me2 (Fig. 4d).

**Fig. 4.**
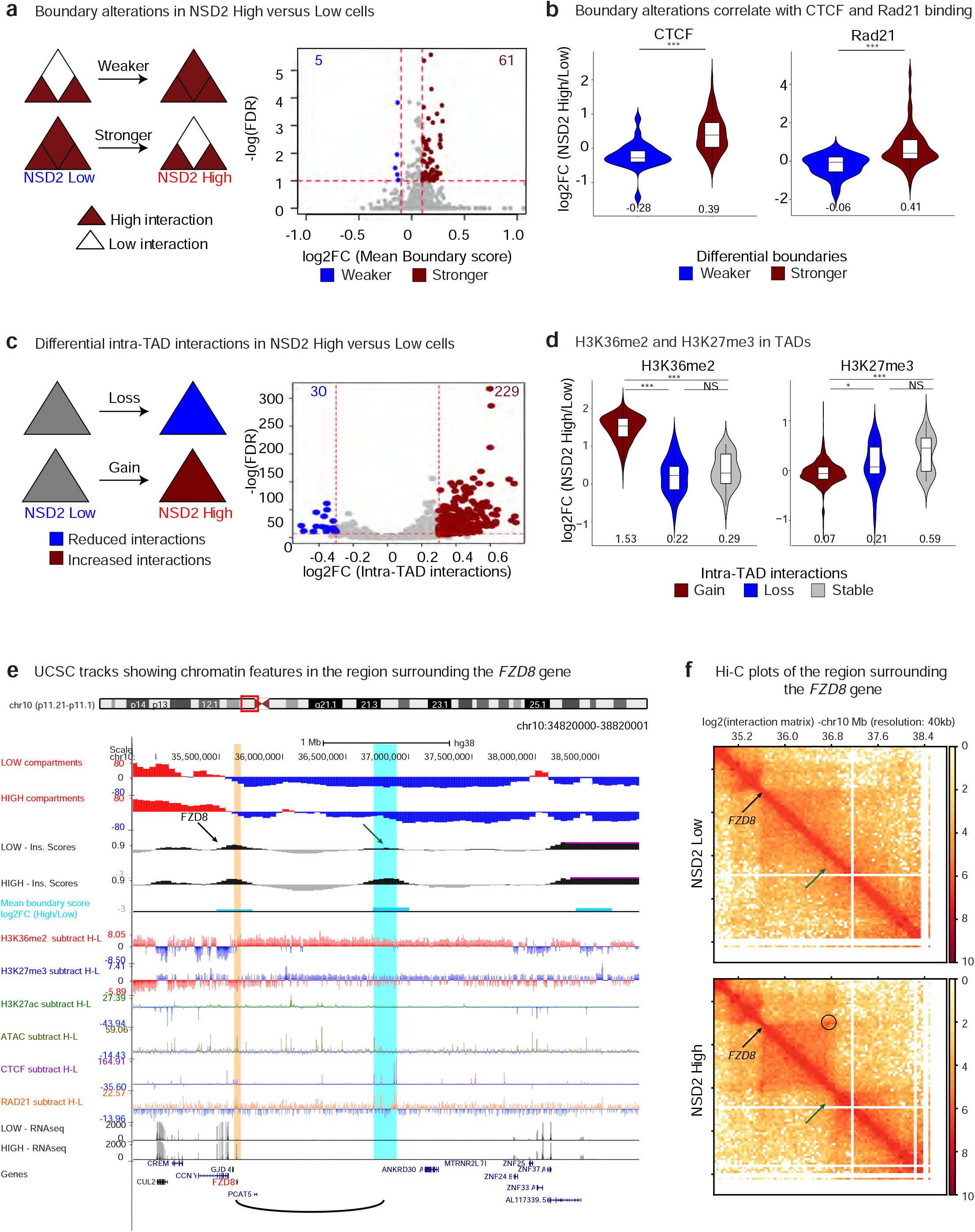
Changes in TAD boundaries and intra-TAD interactions. **a**, Boundary alterations in NSD2 High versus Low cells. NSD2 overexpression is associated with increases (red, 61) and decreases (blue, 5) in TAD boundary strength (cutoffs of absolute Log2 fold change > 0.1 and FDR < 0.05). **b**, TAD boundary increases and decreases in NSD2 High versus Low cells are associated with increases and decreases in CTCF and Rad21 binding, respectively. The median is indicated under the violins. **c**, Intra-TAD interaction changes in NSD2 High versus Low cells for overlapping TADs (1564). NSD2 overexpression is associated with gain (red, 229) and loss (blue, 30) of intra-TAD interactions (cutoffs of absolute Log2 fold change > 0.3 and FDR < 0.05). **d**, Changes in H3K36me2 and H3K27me3 within TADs that have increased (red), decreased (red) or stable (grey) interactions in NSD2 High cells. Log2 Fold changes (NSD2 High/Low cells) for H3K36me2 and H3K27me3 ChIP-seq are shown. The median is indicated under the violins. **e**, UCSC tracks showing chromatin features in the region surrounding the *FZD8* gene (*FZD8* gene indicated in red and location highlighted by a yellow stripe) and the new contact that is formed by a strengthened boundary (blue stripe). The graphical representation of interaction between *FZD8* and the super enhancer is shown as a loop below. H-L refers to subtraction High – Low. **f**, Hi-C plots of the region surrounding the *FZD8* gene. Top panel: NSD2 Low, bottom panel: NSD2 High. Green arrow identifies the TAD boundary that is strengthened in NSD2 High cells versus Low cells. Black arrow indicates the *FZD8* gene. Circle indicates a new loop between *FZD8* and the boundary.

As mentioned above, NSD2 overexpression was associated with a significant strengthening of TAD boundaries concordant with increases in CTCF and Rad21 binding (Fig. 4b). These boundary changes were also associated with transcriptional activation. As an example we highlight activation of *FZD8* (log2 fold-change = 3.16, FDR = 8.18E-34), a gene that is involved in the WNT signaling pathway ^54^ (Fig. 4e). *FZD8* is located at a CTCF enriched boundary that makes a new contact with a downstream region that gains a CTCF/Rad21 site (blue stripe in Fig. 4e) in NSD2 High cells. This is demonstrated by an increase in boundary score (arrows in Fig. 4e, f) and the formation of a new Hi-C loop (circle in Fig. 4f). Consistent with the data in Fig. 2d, the whole domain overlaps a region that is enriched for H3K36me2 and within the new loop there are increases in H3K27ac marks.

Previous studies have proposed that *FGF13* expression enables cells to cope more effectively with RAS-mediated stress and oncogene-mediated excessive protein synthesis, which is known to occur in multiple myeloma in the form of antibody production ^55^. In **Supplementary Fig. 5a, b** we show that activation of *FGF13* occurs at a region with an altered TAD boundary and increased intra-TAD interactions. This region harbors newly formed H3K27ac peaks, enriched chromatin accessibility and overlaps a domain that is enriched for H3K36me2 that undergoes B to A compartment switching. Increased CTCF and Rad21 binding are at the edges of increased Hi-C loops, suggesting that newly formed insulated neighborhoods could facilitate the interaction between *FGF13* and regulatory elements.

We also found preexisting CTCF independent contacts between H3K27ac enriched regions that fall within a loop that has enriched CTCF binding. As an example, we highlight the *KRAS* oncogene (yellow strip **Supplementary Fig. 5c,d**) that plays an important role in driving the multiple myeloma phenotype ^56,57^. As shown in **Supplementary Fig. 5b**, upregulation of *KRAS* (log2 fold change = 0.86, P-value=1.28E-07, FDR=1.38E-06) coincides with a preexisting loop between *KRAS* and a super enhancer whose activity is enriched in NSD2 high cells (see arrow and blue stripe **Supplementary Fig. 5c** and arrows and circle in **Supplementary Fig. 5d**). The figure also highlights an increase in intra-TAD activity. The above examples indicate that significant changes in H3K27ac, CTCF and transcriptional output are frequently clustered together.

### NSD2 overexpression drives clustered chromatin and transcriptional changes in insulated domains

To investigate the connections between alterations in gene expression, CTCF and H3K27ac from a global perspective, we focused our analysis on TADs and CTCF-mediated loops. To identify CTCF-mediated loops, we performed CTCF HiChIP in NSD2 High and Low cells ^58^. Statistical analysis of HiChIP data was carried out using FitHiChIP ^59^ (see Supplementary Information for details). CTCF-mediated interactions were considered significant using an FDR <0.01.

To determine whether significant changes in CTCF, H3K27ac and gene expression in NSD2 High were localized together, we focused on common TADs and common CTCF-mediated loops (see details in Supplementary Information). This allowed us to reliably compare between the NSD2 High versus Low condition. The number of significant changes in CTCF, H3K27ac and gene expression in these common domains is shown in (**Supplementary Fig. 6a, b**). In general, we found that TADs and CTCF loops had a median size of 600kb and 210kb, respectively (**Supplementary Fig. 6c**).

As shown in the scheme in Fig. 5a, we selectively analyzed TADs and CTCF-mediated loops containing at least one CTCF and H3K27ac peak as well as at least one transcriptionally active gene in either the NSD2 Low or High condition. Without any filtering we found that up or down regulation of the three variables was positively correlated in both TADs and CTCF-mediated loops as shown by the red and blue dots, where each dot represents a single TAD or CTCF-mediated loop (**Supplementary Fig. 7a, b**). Positively correlated features are shown by pairwise and three-way log2 fold change comparisons in the 2D and 3D plots, respectively. We also included in our analysis a pairwise and three-way comparison of intra-TAD interaction changes and PC1 changes (see methods for details) to analyze compartment changes within insulated regions that contained alterations in CTCF, H3K27ac and transcriptional output (**Supplementary Fig. 7a**). Again, we found that alterations in intra-TAD interactions and compartment switching were concordant within TADs with alterations in CTCF, H3K27ac and transcriptional output. We next redid the pairwise and three-way comparisons restricting our analyses to only the significant changes in CTCF, H3K27ac and transcriptional output within TADs and CTCF-mediated loops and found that the correlation improved (Fig. 5b, c). We also confirmed that high PC1 mean difference values are associated with regions switching from B to A using the Homer algorithm. This analysis showed that 83% of TADs with B to A switching regions (orange dots) overlap with active features in the 3D graph in common TADs with differential changes in CTCF, H3K27ac as well as RNA (25% of the total positively correlated active TADs) (Fig. 5b). The relationship between significant changes in gene expression and concordant versus discordant alterations in H3K27ac and CTCF in TADs and CTCF-mediated loops is shown in **Supplementary Fig. 7c**. These analyses indicate that the vast majority of changes in CTCF, H3K27ac and transcriptional output are positively correlated.

**Fig. 5.**
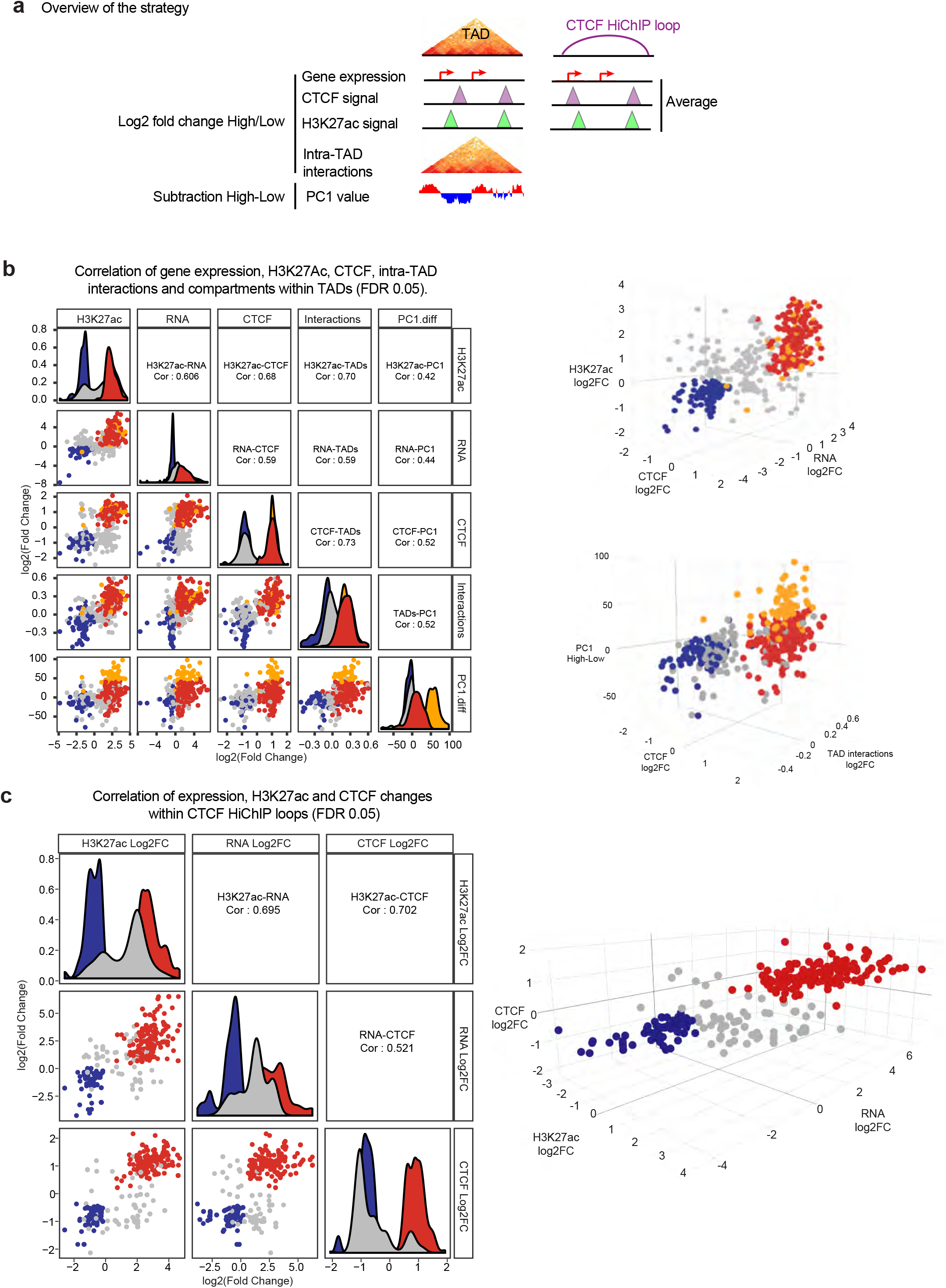
NSD2 overexpression drives concordant chromatin and transcriptional changes in insulated domains. **a**, Scheme illustrating the strategy to identify chromatin and transcriptional changes within TADs or CTCF HiChIP loops that were filtered to have at least one differentially expressed gene, CTCF and H3K27ac peak. **b**, Pairwise (2D scatter plots left panel) and three-way (3D scatter plots right panels) comparisons representing significant log2 fold-changes in gene expression, H3K27ac, CTCF, intra-TAD interactions and PC1 values (representing subtraction of NSD2 High and Low levels) within TADs that have at least one significantly differentially expressed gene, CTCF and H3K27ac peak (FDR < 0.05). Concordant increased and decreased changing TADs are colored in red and blue, respectively. TADs that switch from B to A according to HOMER analysis (see method for details) are highlighted in orange. Pearson correlations are indicated. **c**, Pairwise (2D scatter plots left panel) and three-way (3D scatter plots right panels) comparisons representing significant log2 fold-changes of NSD2 High versus Low levels in gene expression, H3K27ac and CTCF within CTCF HiChIP loops that have at least one differentially expressed gene, CTCF and H3K27ac peak (FDR < 0.05). Concordant increased and decreased changing loops are colored in red and blue, respectively. Pearson correlations are indicated.

### Cells co-opt altered chromatin domains to drive oncogenic transcriptional programs

Within the TADs and loops that harbor significant changes in CTCF, H3K27ac and transcriptional output we detected several newly activated oncogenes associated with multiple myeloma related pathways. These include the protein tyrosine phosphatase gene, *PTPN13* ^60,61^, *FGF13* ^62^ and *ETV5* ^63,64^. The *PTPN13* gene, which regulates cell growth ^65^, differentiation ^60^, mitotic cycle, and oncogenic transformation is located within a CTCF-mediated loop which harbors significant changes in CTCF and H3K27ac (Fig. 6a left), and it is found within a region that undergoes B to A compartment switching. *ETV5*, *a* gene involved in the KRAS pathway ^66,67^, is another example of gene activation that occurs in a CTCF-mediated loop and a TAD that harbors significant changes in CTCF, H3K27ac (Fig. 6a right). As with *PTPN13*, the insulated region contains new H3K27ac and CTCF peaks, overlaps a region that has switched from compartment B to A, and in addition has increased intra-TAD interactions (Fig. 6a right).

**Fig. 6.**
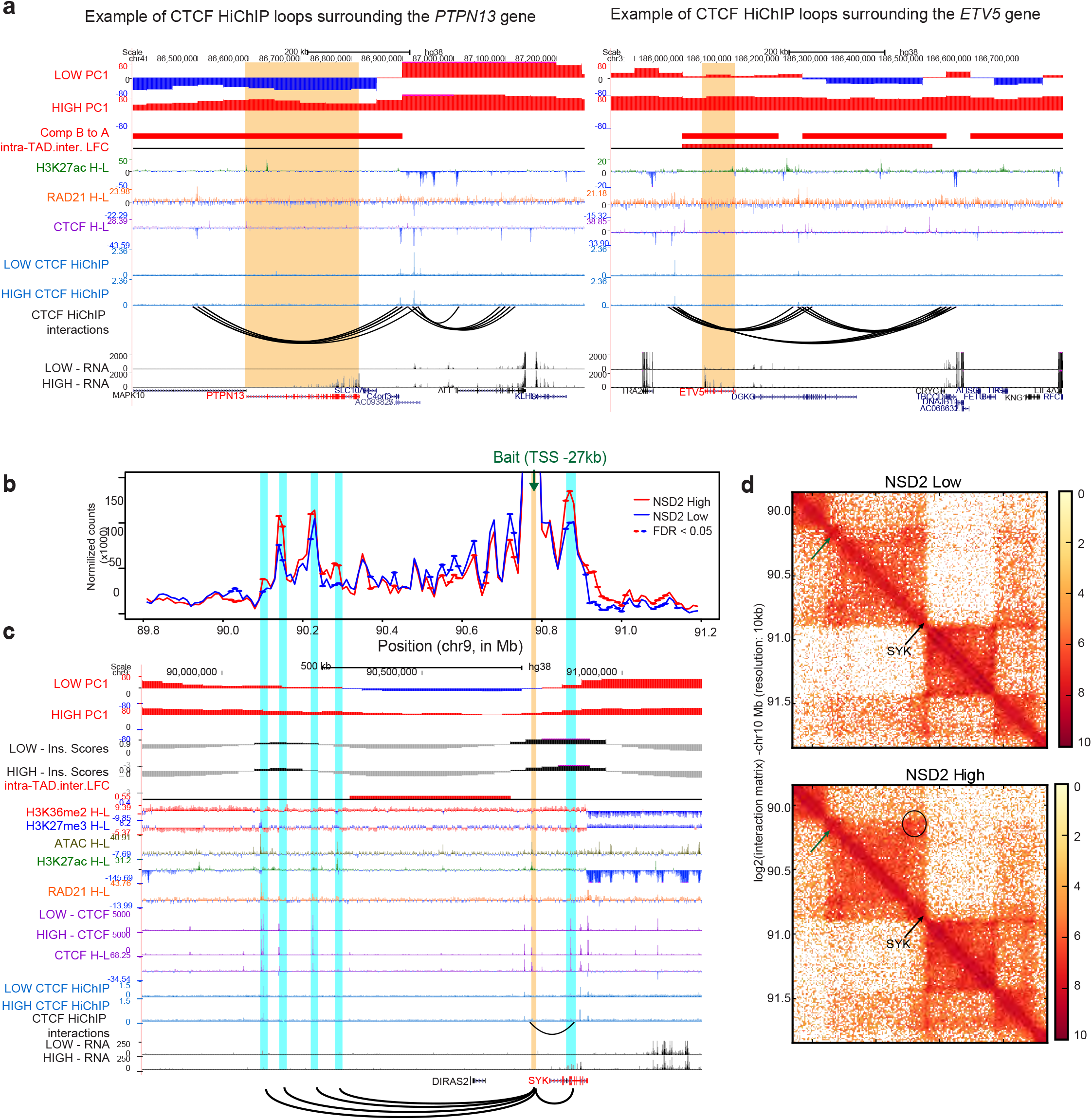
Cells co-opt altered chromatin domains to drive oncogenic transcriptional programs. **a**, UCSC tracks showing chromatin features and CTCF-mediated loop changes in the region surrounding the *PTPN13* (left panel) and ETV5 (right panel) genes (*PTPN13* and ETV5 genes are indicated in red and highlighted with a yellow stripe). H-L refers to subtraction High – Low. **b**, Interaction profile of a 4C bait located 27 kb downstream of the SYK promoter (green arrow, bait −27 kb) in a 2.4 Mb region surrounding the *SYK* gene in NSD2 High (red line represents the average between two replicates) and Low (blue line represents the average between two replicates) cells using 4C counts in 20 kb sliding windows. DESeq2 analysis identified significantly different 4C signal in duplicated 4C samples from NSD2 High versus Low cells. Regions with differential interactions are indicated by red and blue dots (p-value < 0.05). **c**, UCSC tracks showing chromatin features in the region surrounding *SYK* (*SYK* indicated in red). A graphical representation of interactions from the 4C viewpoint (highlighted by a yellow strip) located 27 kb downstream of the *SYK* promoter is drawn with arcs at the bottom and highlighted by blue stripes. H-L refers to subtraction High – Low. **d**, Hi-C plots of the region surrounding *SYK* in NSD2 Low and High cells (left and right panels, respectively).

*SYK* is another example of an oncogene that is activated within a CTCF-mediated loop that has significant increases in H3K27ac and CTCF peaks. To investigate whether upregulation of *SYK* in NSD2 High cells involves alterations in promoter contacts, we performed high resolution 4C-seq from a viewpoint (bait) located on a CTCF site 27Kb downstream of the *SYK* promoter. We identified significantly increased interactions with upstream and downstream regions (FDR < 0.05; marked by the small ovals on the 4C plot in Fig. 6b). All of these sites correspond to enriched CTCF or H3K27ac peaks that overlap with enriched cohesin peaks in the NSD2 High cells (Fig. 6c). The interactions identified by 4C-seq can also be visualized by Hi-C (arrows and circles in Fig. 6d). Interestingly, the change in interactions seen in Fig. 6b overlap with a region that changes from compartment B to A and has increased intra-TAD interactions. As in other examples, these changes all occur in a domain where H3K36me2 is enriched in the NSD2 High cells. Taken together our data indicate that gene expression changes and activation of oncogenes in NSD2 overexpressing cells are found in insulated regions with accompanying alterations in H3K27ac and CTCF peaks.

### Changes in gene expression occur as a function of changes of chromatin in TADs and CTCF loops

To better understand the relationship between transcriptional changes and chromatin changes in TADs and CTCF loops we modeled the probability that a gene is differentially expressed, as a function of changes in CTCF and/or H3K27ac within these insulated domains, using logistical regression. First, we assigned all genes, CTCF, and H3K27ac peaks to all TADs and CTCF loops in the NSD2 High and Low conditions. Then, we aggregated these and computed whether a gene shares a TAD or CTCF loop with a differential CTCF and/or H3K27ac peak. Finally, we checked whether changes in CTCF or H3K27ac peaks between the two conditions in the same TAD or CTCF loop have an effect on whether a gene is differentially expressed (Fig. 7a). We found that changes in H3K27ac significantly increase the probability that a gene sharing the same CTCF loop is differentially expressed, while changes in both CTCF and/or H3K27ac peaks significantly increase the probability that a gene sharing the same TAD is differentially expressed. These changes represent 60.4% of the differentially expressed genes. Changes in gene expression cluster with changes in CTCF and/or H3K27ac in only a subset of TADs (17%) and loops (13.8%). The number of CTCF and H3K27ac changes that occur within TADs and loops that have altered gene expression is shown in Fig. 7b.

**Fig. 7.**
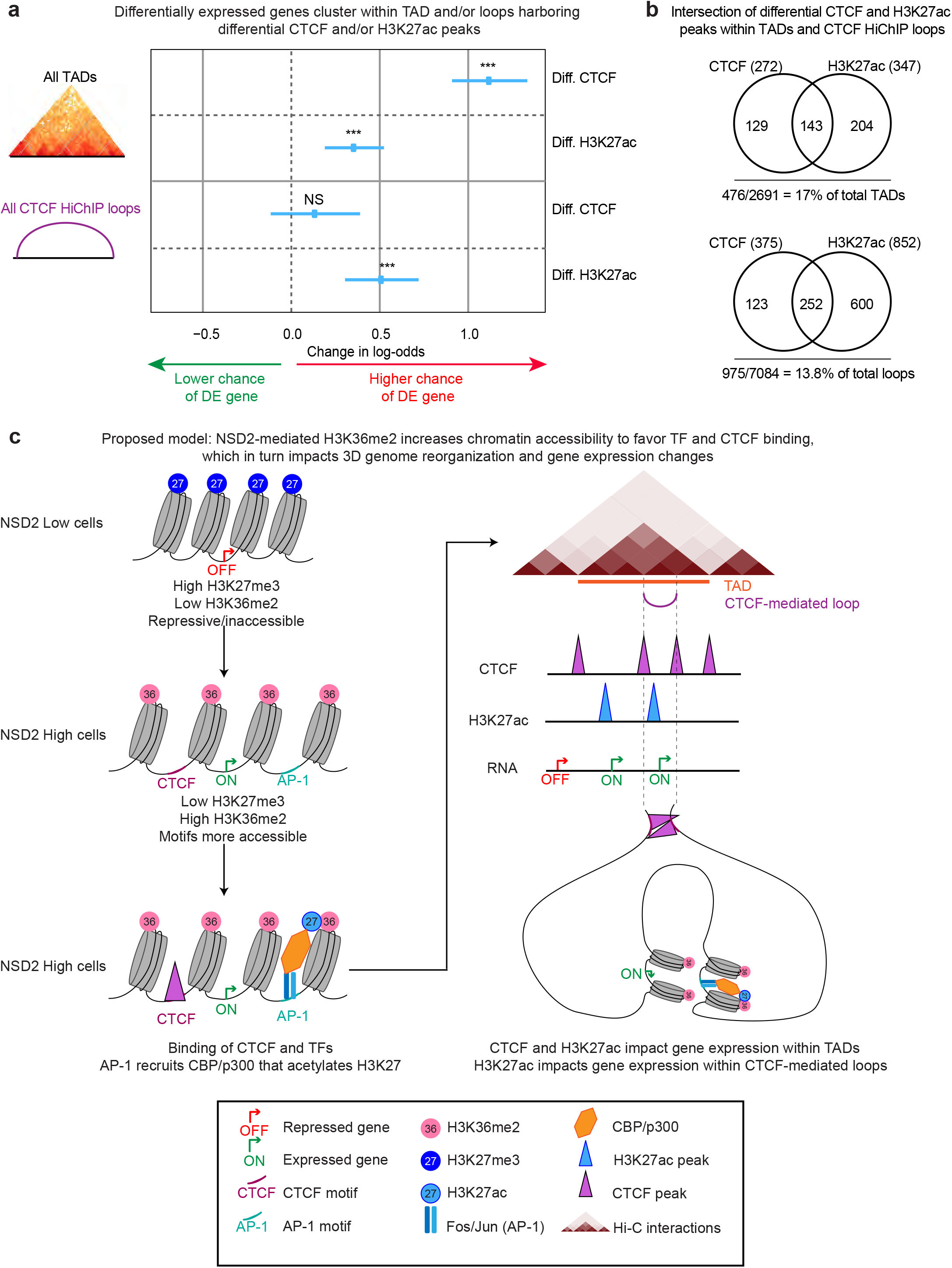
Changes in gene expression occur as a function of changes in chromatin in TADs and CTCF loops. **a**, Logistical regression model of gene expression changes as a function of CTCF and/or H3K27ac changes in TADs and CTCF loops (see supplementary information for method details). Diff. CTCF and H3K27ac refer to differential CTCF and H3K27ac peaks. A DE gene refers to a differentially expressed gene. P-values were calculated using Wald-test (\*\*\**P*< 2e-16 for Diff. CTCF in TADs; \*\*\**P*=2.6e-5 for Diff. H3K27ac in TADs; ^NS^*P*= 0.29 for Diff. CTCF in loops; \*\*\**P*=1.38e-6 for Diff. H3K27ac in loops). **b**, Intersection of differential CTCF and H3K27ac peaks within TADs and CTCF loops. **c**, Proposed model: NSD2-mediated H3K36me2 increases chromatin accessibility to favor TF and CTCF binding, which in turn impacts 3D genome reorganization and gene expression changes.

## Discussion

Deletion of individual CTCF sites is known to disrupt chromosome interactions allowing the spreading of active chromatin and altering gene regulation ^19–21^. However, it is not known whether alterations in chromatin can affect CTCF binding, 3D genome organization and gene expression. To address this question we examined the effect of NSD2 overexpression in t(4;14) multiple myeloma. The activation of oncogenic transcriptional pathways in this context has been shown to rely on the histone methyl transferase activity of NSD2 and deposition of H3K36me2 ^33^. Here we show that NSD2 overexpressing cells can coopt altered chromatin domains to drives changes in 3D organization including A/B compartmentalization and TAD architecture, linked to the activation of oncogenic transcriptional pathways.

Expansion of H3K36me2 outside of active gene bodies provides a favorable environment for enrichment of H3K27ac and binding of CTCF. Hi-C and HiChIP analyses reveal that changes in H3K27ac, CTCF and transcriptional activity occur in a predominantly concordant manner in common TADs (72%) and CTCF loops (82%). Furthermore, intra-TAD interactions and B to A compartment changes are positively correlated with changes in H3K27ac, CTCF and transcriptional activity in TADs. Importantly, we find that 60.4% of the deregulated genes cluster together in a subset of TADs (17%) or CTCF-mediated loops (13.8%) that harbor significant changes in CTCF and/or H3K27ac binding. Using a linear regression model, we demonstrate that changes in H3K27ac significantly increase the probability that a gene sharing the same CTCF loop is differentially expressed, while changes in both CTCF and/or H3K27ac peaks significantly increase the probability that a gene sharing the same TAD is differentially expressed. Our findings are consistent with a model in which gene regulation is constrained at the 3D level by insulated boundaries that restrict the influence of enhancers. As highlighted in the examples in this manuscript, these changes provide an explanation for the activation of many oncogenes associated with multiple myeloma.

What is the relationship between alterations in H3K36me2, H3K27ac, CTCF and A/B compartment changes? Since compartment switching is linked to changes in chromatin activity it is possible that changes in H3K36me2 on its own could induce compartment changes. It would be difficult to tease this apart because spreading of H3K36me2 into intergenic regions is accompanied by changes in transcriptional activity, H3K27ac, CTCF, ATAC-seq and Rad21, all of which could contribute to changes in compartmentalization. We speculate however that it is the overall increases in CTCF/Rad 21 binding that plays a key role in compartment weakening, in line with the finding that acute depletion of cohesin and loss of TAD structure leads to strengthening of compartments ^9,10^. Thus, although compartments and TADs are formed through independent pathways, TAD structures interfere with compartmentalization.

Since the NSD2 phenotype relies on the histone methyl transferase activity of the protein, the upstream causal event is the expansion of H3K36me2 domains. Beyond this change it is not clear what order the other events occur in, or whether they are reliant on each other. The causal directionality would be difficult to tease apart as we could not exclude the possibility that chromatin changes are inter-dependent. It is unlikely that alterations in H3K27ac enrichment and CTCF binding are dependent on each other as they do not bind to overlapping sites. We speculate that opening up of the chromatin through deposition of the active mark, H3K36me2 allows binding of transcription factors such as AP-1, which recruits CBP/p300 ^68^ that mediates deposition of H3K27ac, a well-defined marker of enhancer activity (Fig. 7c). Indeed, our studies reveal that the motif for AP-1 is enriched at new H3K27ac peaks, including those designated as super-enhancers and moreover expression of *FOS2L* and *JUN* that encode the AP-1 subunits are upregulated in NSD2 high cells. AP-1 is enriched in activation hubs that connect enhancers and promoters in macrophages ^69^, and it may participate in regulating enhancer-promoter contacts in NSD2 overexpressing multiple myeloma cells. Similarly, we speculate that H3K36me2-mediated accessibility promotes binding of CTCF. H3K27ac enriched regulatory elements and CTCF bound sites are known to participate in cohesin-mediated loop formation and in line with this, we show that new H3K27ac and CTCF peaks overlap with new Rad21 peaks and form new or strengthened contacts linked to changes in gene expression. An overall increase in CTCF/Rad21 and H3K27ac as well as transcriptional activity likely contributes to the overall increase in intra-TAD interactions as shown by the pairwise and three way-comparisons in Fig. 5. These changes are also positively correlated with B to A compartment changes. Specifically, we find that 85% of regions that switch from B to A are found in TADs with a significant increase in CTCF, H3K27ac and transcriptional output and perhaps not surprisingly compartment switching occurs in TADs with the highest activity of all three.

Here we have identified a functional interplay between NSD2-mediated changes in chromatin, 3D organization and transcriptional output. These results highlight the bidirectional relationship between 2D chromatin status and 3D genome organization in gene regulation. In the context of NSD2 overexpression in multiple myeloma, we demonstrate that alterations in CTCF and H3K27ac increase the probability of changing gene expression within the same TAD or CTCF loop. These chromatin changes are associated with the majority of changes in gene expression including genes such as *KRAS* and *SYK*, that in the absence of driver mutations contribute to the oncogenic program of the disease as highlighted by the enrichment of KRAS pathway genes amongst deregulated genes (**Supplementary Fig. 1b**). This work demonstrates that alterations in chromatin modifiers in disease states enables dissection of the interplay between 2D and 3D chromatin structure and their links to gene regulation.

## Methods

Methods are available in the note section of supplementary information.

## Acknowledgements

The authors thank the Ben Ho Park laboratory for sending the NSD2 High and Low cells, Skok lab members for helpful scientific discussions, New York University School of Medicine High Performance Computing (HPC) for computing technical support, and Adriana Heguy and the Genome Technology Center (GTC) core for sequencing efforts.

## Author contributions

Conceptualization & Study Design, P.L and J.S; Investigation, P.L; Formal analysis, P.L, Sa.Ba, J-R.H, T.S, G.S, A.K, M.C, So.Bh; Writing – Original Draft, P.L and J.S; Supervision, R.B, F.A, A.T and J.S; Funding Acquisition, P.L, J.S, A.T.

