## Supplementary Figures ad Methods for "NSD2 overexpression drives clustered chromatin and transcriptional changes in a subset of insulated domains"

[1]Dept. of Pathology, New York University Langone Health, New York, NY 10016, USA; [2]Dept. of Biology, Center for Genomics and Systems Biology, NYU, New York, NY 10003, USA; [3]Division of Vaccine Discovery, La Jolla Institute for Immunology, La Jolla, CA 92037, USA; [4]School of Medicine, University of California, San Diego, La Jolla, CA 92093, USA; [5]Applied Bioinformatics Laboratories, NYU School of Medicine, New York, NY 10016, USA. [6]Laura and Isaac Perlmutter Cancer Center, NYU School of Medicine, New York, NY 10016, USA.

* Equal contribution.

[#] Corresponding author

Running title: NSD2 overexpression drives 3D genome reorganization

#### SUPPLEMENTARY FIGURES

**Supplementary Figure 1. RNAseq analysis in NSD2 High versus Low cells**. **a,** PCA of RNA-seq replicates for NSD2 High and Low cells. **b,** Gene Set Enrichment Analysis of gene expression in NSD2 High versus Low cells on normalized genes reads counts for each replicate.

**Supplementary Figure 2. Transcription factor motifs identified in differential H3K27ac peaks**. **a,** PCA of H3K27ac ChIP-seq replicates for NSD2 High and Low cells. **b,** Identification of super-enhancers in NSD2 Low (upper panel) and High (lower panel) using H3K27ac with 'ROSE' (rank ordering of super-enhancers [1]). **c,** Transcription factor motifs identified in increased (1650) and decreased (303) H3K27ac peaks using TRAP, related to Figure 1. Motifs also presents in Figure 1G are in bold. Motifs found in increased (red) and decreased (blue) H3K27ac peaks (-log10 p-value). **d,** PCA of CTCF ChIP-seq replicates for NSD2 High and Low cells.

**Supplementary Figure 3. Chromatin landscape of the regions surrounding CTCF and H3K27ac changes**. Heatmaps **a,** and average profiles **b,** of H3K27ac, H3K36me2, H3K27me3, ATAC-seq, Rad21 and CTCF signal at the top 1000 increased, 100 stable and 1000 decreased H3K27ac peaks in NSD2 High versus Low cells. Heatmaps **c,** and average profiles (`D) of CTCF, H3K36me2, H3K27me3, ATAC-seq, Rad21 and H3K27ac signal at top 1000 increased, 100 stable and 1000 decreased CTCF peaks in NSD2 High versus Low cells. Top 1000 increased, 100 stable and 1000 decreased CTCF and H3K27ac peaks are based on Log2 Fold changes in reads counts in NSD2 High versus Low cells. Peaks are ranked by H3K27ac **a,** and CTCF **c,** signal in NSD2 High cells.

**Supplementary Figure 4. Hi-C analysis in NSD2 High and Low cells. a,** Histogram representing read filtering of Hi-C replicates processed using Hi-C bench [2]. "Ds accepted-intra" and “ds accepted inter” reads were retained for further analysis. **b,** PCA of Hi-C replicates for NSD2 High and Low cells processed with Hi-C bench at a resolution of 40 kb. Histograms representing the total size **c,** and average size **d,** of A, B and switching compartments in NSD2 High and Low cells. **e,** Bar plot representing the proportion of A (red) and B (blue) compartments in the genome for NSD2 High and Low cells. **f,** Histogram representing the number of TADs for each replicate of NSD2 High and Low cells. **g,** Boxplots representing the distribution of TAD sizes (in megabases, Mb), for each replicate of NSD2 High and Low cells.

**Supplementary Figure 5. a,** UCSC tracks showing chromatin features in the region surrounding the *FGF13* gene (*FGF13* gene indicated in red and location highlighted by a yellow stripe) and the new contacts that are formed by stronger CTCF and Rad21 binding (blue stripes and shown as a loop below). H-L refers to subtraction High – Low. **b,** Hi-C plots of the region surrounding the *FGF13* gene. Top panel: NSD2 Low, bottom panel: NSD2 High. Black arrow indicates the *FGF13* gene. Circle indicates a loop between *FGF13* and surrounding increased CTCF and Rad21 regions highlighted by a blue strip in panel **b,**. **c,** UCSC tracks showing chromatin features in the region surrounding the *KRAS* gene (*KRAS* gene indicated in red and highlighted by a yellow stripe) and a downstream super enhancer (indicated by a green arrow and highlighted by a blue stripe). Left and right boundaries surrounding intra-TAD interaction gain are highlighted by blue stripes and indicated by green arrows labelled as boundary ‘1’ and ‘2’, respectively. A graphical representation of the interaction between *KRAS* and the super enhancer is shown as a loop below. H-L refers to subtraction High – Low. **d,** Hi-C plots of the region surrounding the *KRAS* gene in NSD2 Low versus High cells (top and bottom panels, respectively). Black arrow indicates the location of *KRAS*. Green arrows indicate the super enhancer (also labelled as “SE”), and left and right boundaries surrounding intra-TAD interaction gain as labelled as boundary ‘1’ and ‘2’, respectively. Circle indicates a loop between *KRAS* and the super enhancer.

**Supplementary Figure 6. NSD2 overexpression drives concordant chromatin and transcriptional changes in insulated domains. a,** Bar plot (left panel) and Volcano plots (right panel) showing significant NSD2-mediated changes in CTCF binding (left panels), gene expression (middle panels) and H3K27ac signal (right panels) within TADs (top panels) or CTCF HiChIP loops (bottom panels). Increases of log2 fold change >1 are shown in red and decreases, log2 fold change <-1 are shown in blue (FDR <0.01). **b,** Percentage of significantly differential features in TADs. **c,** Density plot of CTCF HiChIP loop sizes. **c,** Bar graph showing the proportion of concordant and discordant change in CTCF, H3K27ac and gene expression in TADs and CTCF loops.

**Supplementary Figure 7. NSD2 overexpression drives concordant chromatin and transcriptional changes in insulated domains. a,** Pairwise (2D scatter plots left panel) and three-way (3D scatter plots right panels) comparisons representing log2 fold-changes of gene expression, H3K27ac, CTCF, intra-TAD interactions and PC1 values (representing subtraction of NSD2 High and Low levels within TADs) in NSD2 High versus Low cells. Concordant increased and decreased changing TADs are colored in red and blue, respectively. TADs that switch from B to A as determined by HOMER analysis (see method for details) are highlighted in orange. Pearson correlations are indicated. **b,** Pairwise (2D scatter plots left panel) and three-way (3D scatter plots right panels) comparisons representing log2 fold-changes of gene expression, H3K27ac and CTCF within CTCF HiChIP loops in NSD2 High versus Low cells. Concordant increased and decreased changing loops are colored in red and blue, respectively. Pearson correlations are indicated.

#### SUPPLEMENTARY NOTE

#### SUPPLEMENTARY METHODS

##### Cell lines

NSD2 High (one clone of NTKO and KMS11 parental cell line, which is a human myeloma cell line from a female patient) and NSD2 Low cells (two clones of TKO) were obtained from Ben Ho Park [3] who generated the cell lines. Upon reception, the phenotype of the cells was as described [3], and NSD2 High and Low cells harbored expected alterations in NSD2 RNA and protein as confirmed by RNA-seq and Western-blot, respectively. Additional cell authentication was not performed. Cells were maintained in culture as per Ben Ho Park's recommendations. Briefly, cells were grown in RPMI-1640 supplemented with 10% fetal bovine serum, 100 U/mL penicillin and 100 µg/mL streptomycin and subcultured twice a week by dilution. Our PCA revealed that NTKO (two replicates from independent cultures) and KMS11 (one replicate) were identical in terms of gene expression, CTCF and H3K27ac peaks. Only NTKO cells were used for further experiments. For NSD2 low cells, two replicates from independent cultures were used from one clone as well as one replicate from the other clone. Our PCA revealed that TKO clones and replicates were identical in term of gene expression, CTCF and H3K27ac peaks. For downstream experiments, NTKO and KMS11 cells were trypsinized while TKO cells were directly obtained from suspension cultures. Cells were freshly crosslinked for ChIP-seq, Hi-C, Hi-ChIP and 4C (see corresponding sections for detailed protocols), resuspended in RLT buffer for RNA extraction as per the instructions provided by the kit (RNeasy plus kit from QIAGEN), or directly processed for ATAC-seq (see corresponding section for detailed protocol).

**RNA-seq**

RNA was extracted from 5 replicates for each condition, using the RNeasy plus kit from QIAGEN. Poly-adenylated transcripts were positively selected using the NEBNext® Poly**a,** mRNA Magnetic Isolation Module following the kit procedure. Libraries were prepared according to the directional RNA-seq dUTP method adapted from http://wasp.einstein.yu.edu/index.php/Protocol:directional WholeTranscript_seq that preserves information about transcriptional direction. Sequencing was performed with Illumina Hi-Seq 2500 using 50 cycles paired-end mode.

**ATAC-seq**

NTKO and KMS11 cells were trypsinized while TKO cells were directly obtained from suspension cultures. Cell were counted to collect 50,000 cells per replicate. Two different cultures of NTKO cells were used for the NSD2 High condition and one culture from two different clones of TKO for the NSD2 Low condition. The procedure was repeated in three independent days for a total of six replicates for NSD2 High and six replicates for NSD2 Low. The assay was performed as described previously [4]. Cells were washed in cold PBS and resuspended in 50 µl of cold lysis buffer (10 mM Tris-HCl, pH 7.4, 10 mM NaCl, 3mM MgCl2, 0.1% IGEPAL CA-630). The tagmentation reaction was performed in 25 µl of TD buffer (Illumina Cat #FC-121-1030), 2.5 µl Nextera Tn5 Transposase, and 22.5 µl of Nuclease Free H2O at 37$^{o}$C for 30 min. DNA was purified on a column with the Qiagen Mini Elute kit, eluted in 10  µl H2O. Purified DNA (10 µl) was combined with 10 µl of H2O, 2.5 µl of each primer at 25 mM and 25 µl of NEB Next PCR master mix. DNA was amplified for 5 cycles and a monitored quantitative PCR was performed to determine the number of extra cycles needed as per the original ATAC-seq protocol [4]. DNA was purified on a column with the Qiagen Mini Elute kit. Samples were quantified using Tapestation bioanalyzer (Agilent) and the KAPA Library Quantification Kit and sequenced on the Illumina Hi-Seq 2000 using 50 cycles paired-end mode. The six replicates were sequenced independently and three replicates were pooled together during the data processing in order to get two replicates with sufficient sequencing depth for downstream analysis (see ATAC-seq processing data for details).

**ChIPmentation**

Cells were fixed in culture medium within 1% formaldehyde at RT for 10 minutes and quenched with 0.125 M glycine. Pellets were washed twice with ice-cold PBS, snap-frozen in liquid nitrogen and stored at -80°C. ChIP-seq was performed as per the original ChIPmentation protocol [5] in triplicate for CTCF and H3K27ac, and in duplicate for H3K36me2 and Rad21. Briefly, chromatin was lysed during a 10 min rotation in the cold room in 350 µl of lysis buffer (10 mM Tris-HCl pH 8.0, 100 mM NaCl, 1 mM EDTA pH8.0 NaOH, 0.5 mM EGTA pH8.0 NaOH, 0.1% sodium deoxycholate, 0.5% N-lauroysarcosine). Lysates were sonicated using Bioruptor (Diagenode) (15 cycles 30 sec ON, 30 sec OFF, an agarose gel was run to make sure that the sonicated DNA smear was in the range of 100-700bp).Triton X-100 1% finale were added and the samples were centrifuged 5 min at 16000 rcf at 4°C. Supernatant was collected. Antibody was combined with protein A magnetic beads for one hour at room temperature and added to chromatin. For CTCF, H3K27ac and Rad21 and IgG as negative control, 10 µl of antibody (Millipore 07-729, Abcam ab4729, Abcam ab992, Abcam ab37415, respectively) was added to 50 µl of protein-A magnetic beads (Dynabeads) and added to the sonicated chromatin from 10 million cells per immunoprecipitation. For H3K36me2, internal spike-in was added for normalization as previously described [6]. Briefly, per immunoprecipitation, 1 µl of H3K36me2 antibody and 0.1 µl of Drosophila-specific H2Av antibody were added to 10 µl of protein-A magnetic beads and added to 100 µg of human sonicated chromatin supplemented with 2 µg of drosophila sonicated chromatin. Of note, chromatin was quantified with Nanodrop at 260 nm. Immunoprecipitation was performed for 3 to 6 hours rotating in the cold room, then washes and tagmentation were performed as per the original ChIPmentation protocol [5]. Briefly, beads were washed twice with 500 µl cold low-salt wash buffer (20 mM Tris-HCl pH 7.5, 150 mM NaCl, 2 mM EDTA pH8.0 NaOH, 0.1% SDS, 1% triton

X-100), twice with 500 µl cold LiCl-containing wash buffer (10 mM Tris-HCl pH 8.0, 250 mM LiCl, 1 mM EDTA pH8.0 NaOH, 1% triton X-100, 0.7% sodium deoxycholate), and twice with 500 µl cold 10 mM cold Tris-Cl, pH 8.0, to remove detergent, salts and EDTA. Subsequently, beads were resuspended in 25 µl of the freshly prepared tagmentation reaction buffer (10 mM Tris-HCl, pH 8.0, 5 mM MgCl2, 10% dimethylformamide) and 1 µl Tagment DNA Enzyme from the Nextera DNA Sample Prep Kit (Illumina) and incubated at 37°C for 1 min in a thermocycler. Following tagmentation, the beads were washed twice with 500 µl cold low-salt wash buffer (20 mM Tris-HCl pH 7.5, 150 mM NaCl, 2 mM EDTA pH8.0 NaOH, 0.1% SDS, 1% triton X-100), and twice with 500 µl cold Tris-EDTA-Tween buffer (0.2% tween, 10 mM Tris-HCl pH 8.0, 1 mM EDTA pH 8.0). Chromatin was eluted and decrosslinked by adding 70 µl of freshly prepared elution buffer (0.5% SDS, 300 mM NaCl, 5 mM EDTA pH 8.0, 10 mM Tris-HCl pH 8.0) and 2 µl of proteinase K at 10 mg/ml for 2 hours at 55°C and overnight incubation at 65°C. Supernatant was kept and to recover as much DNA as possible, beads were washed with an additional 30 µl of elution buffer and combined supernatant was incubated an additional hour at 55°C. DNA was purified on a column with the Qiagen Mini Elute kit. Purified DNA (20 µl) was combined with 2.5 µl of each primer at 25 mM and 25 µl of NEB Next PCR master mix and was amplified as per the ChIPmentation protocol (Schmidl et al. 2015) in a thermomixer with the following program: 72°C for 5 min; 98°C for 30 s; 14 cycles of 98°C for 10 s, 63°C for 30 s and 72°C 30 s; and a final elongation at 72°C for 1 min. DNA was purified using two consecutive rounds of SPRI AMPure XP beads: the first one with a beads-to-sample ratio of 0.6:1 to remove potential fragments larger than 700 bp (supernatant kept) and the second one with a beads-to-sample ratio of 1:1 to remove potential primer dimers (beads kept), and eluted in 20 µl of H2O. Samples were quantified using Tapestation bioanalyzer (Agilent) and KAPA Library Quantification Kit and sequenced with Illumina Hi-Seq 2500 using 50 cycles paired-end mode (CTCF and H3K27ac) or single-end mode (Rad21 and H3K36me2).

**Hi-C**

Hi-C was performed in duplicate, from 0.5 to 1 million cells fixed in culture medium within 1% formaldehyde at RT for 10 minutes and quenched with 0.125M glycine. Hi-C samples were processed using the Arima Hi-C kit as per the kit protocol, and sequenced with Illumina Hi-Seq 2500 using 50 cycles paired-end mode.

**CTCF HiChIP**

HiChIP was performed in duplicate. Cells were fixed in culture medium with 1% formaldehyde at RT for 10 minutes and quenched with 0.125 M glycine. Pellets were washed twice with ice-cold PBS, snap-frozen in liquid nitrogen and stored at -80°C. HiChIP was performed with 15 million cells, as per the original protocol [7]. Cells were then lysed in 500 µl ice-cold lysis buffer (10 mM Tris-HCl pH 8.0, 10 mM NaCl, 0.2% Igepal CA-630, protease inhibitor cocktail (Roche complete, EDTA-free)) rotating at 4°C for 30 minutes. Cell pellets were collected, washed once in 500 µl ice-cold lysis buffer and then incubated in 100 µl 0.5% SDS at 62°C for 10 min. SDS was quenched by adding 285 µl of H2O and 50 µl of Triton X-100 10%, and incubating at 37°C for 15 min. Chromatin was then digested by adding 50 µl of NEBuffer 2 10X and 350 units of MboI (NEB R0147M) at 37°C for 2 hours while rotating at 950 rpm. MboI was inactivated by incubating the samples 20 minutes at 62°C. To fill in the restriction fragment overhangs and mark the DNA ends with biotin, 1.5µl 10 mM dCTP, 1.5µl 10 mM dGTP, 1.5µl 10 mM dTTP, 37.5µl 0.4 mM biotin-14-dATP (Life Technologies 19524-016), and 10 µL 5U/µl Klenow (DNA polymerase I large fragment, NEB M0210L) were added to each tube, and incubated for 60 minutes at 37°C. Ligation mix was added to the samples (150 µl 10x ligation buffer (NEB B0202S), 7.5 µl 20mg/ml BSA (NEB B9001S), 150 µl Triton X-100 10%, 10 µl 400U/µl T4 DNA ligase (NEB M0202S), and 655.5 µl H2O) for 4 hours at RT with rotation. Following ligation, nuclei were pelleted and resuspended in 350 µl cold Nuclei Lysis Buffer (50 mM Tris-HCl pH 7.5, 10 mM EDTA, 1% SDS, and 1x Protease Inhibitors) with incubation rotating in the cold room for 10 min. Samples were sonicated on the bioruptor for 15 min (an agarose gel was performed to make sure that the sonicated DNA smear was in the range of 250-600bp), supplemented with 1% Triton X-100 and centrifuged 5 min at 16000 rcf at 4°C. CTCF antibody (5 µl, Millipore 07-729) was combined with 50 µl of protein-A magnetic beads (Dynabeads) for one hour at room temperature, and added to sonicated chromatin from 15 million cells. Immunoprecipitation was performed by overnight incubation rotating in cold room and washes were performed. Briefly, beads were washed twice with 500 µl cold Low-salt wash buffer (20 mM Tris-HCl pH 7.5, 150 mM NaCl, 2 mM EDTA pH8.0 NaOH, 0.1% SDS, 1% triton X-100), twice with 500 µl cold high-salt wash buffer (20 mM Tris-HCl pH 7.5, 500 mM NaCl, 2 mM EDTA pH8.0 NaOH, 0.1% SDS, 1% triton X-100), and twice with 500 µl cold LiCl-containing wash buffer (10 mM Tris-HCl pH 8.0, 250 mM LiCl, 1 mM EDTA pH8.0 NaOH, 1% NP-40, 1% sodium deoxycholate). Chromatin was eluted and decrosslinked by adding 100 µl of freshly prepared elution buffer (0.5% SDS, 300 mM NaCl, 5 mM EDTA pH 8.0, 10 mM Tris-HCl pH 8.0) and 10 µl of proteinase K at 10 mg/ml for 45 min at 55°C and at least 1.5 hour at 67°C. DNA was purified on kept a column with the Qiagen Mini Elute kit, eluted in 12 µl of H2O, and quantified using Qubit. Of note, we obtained between 3 to 8 ng of DNA for CTCF HiChIP from 15 million cells. To enrich for ligation events, 5 µl of Streptavidin C-1 beads were washed in Tween Wash Buffer (TWB, 5 mM Tris-HCl pH 7.5, 0.5 mM EDTA pH 8.0, 1M NaCl, 0.05% Tween-20), resuspended in 10 µl of 2X Biotin Binding Buffer (10 mM Tris-HCl pH 7.5, 1 mM EDTA pH 8.0, 2 M NaCl), added to 10 µl of the samples and incubated at room temperature for 15 min with rotation. Samples were then washed twice in TWB with 2 min incubation at 55°C shaking. For tagmentation, beads were washed twice in 100 µl of freshly prepared tagmentation reaction buffer (10 mM Tris-HCl, pH 8.0, 5 mM MgCl2, 10% dimethylformamide), and resuspended in 25 µl of the tagmentation reaction buffer and 1 µl Tagment DNA Enzyme from the Nextera DNA Sample Prep Kit (Illumina) and incubated at 55°C for 10 min in a thermocycler with interval shaking. Beads were resuspended in 50 mM EDTA and incubated at 50°C for 30 min to quench the transposase reaction. This was followed by two washes in 50 mM EDTA incubated at 50°C for 3 min, two washes in Tween Wash Buffer incubated at 55°C for 2 min, and one wash in 10 mM Tris-HCl pH 7.5. Beads were resuspended in 50 µl of PCR master mix (1 µl of each primer at 25 mM, 25 µl of NEB Next PCR master mix and 23 µl of H2O) and DNA was amplified in a thermomixer with the following program: 72°C for 5 min; 98°C for 30 s; 10 cycles of 98°C for 10 s, 63°C for 30 s and 72°C for 1 min. DNA was purified using two consecutive rounds of SPRI AMPure XP beads: the first one with a beads-to-sample ratio of 0.6:1 to remove potential fragments larger than 700 bp (supernatant kept) and the second one with a beads-to-sample ratio of 0.18:1 to keep fragments greater than 300 bp (on beads), and eluted in 15 µl of H2O. Samples were quantified using the Tapestation bioanalyzer (Agilent) and the KAPA Library Quantification Kit and sequenced with Illumina Hi-Seq 2500 using 50 cycles paired-end mode.

**4C-seq**

4C-seq was performed in duplicate and analyzed as previously described [8,9] with minor changes. 10 million cells were fixed in 2% formaldehyde for 10 minutes at room temperature and quenched with glycine (0.125 M final concentration). Nuclei were isolated in lysis buffer (50mM Tris-HCl ph7.5, 150mM NaCl, 5mM EDTA, 0.5% NP-40, 1% TX-100 containing 1X Roche complete Mini protease inhibitors) and dounced 40 times on ice. Nuclei were resuspended in 360 µl H2O and 60 µl warm 10X NEB DpnII buffer. They were permeabilized using 15 µl 10% SDS for 60min at 37°C and then 150 µl 10% Triton X-100 for 60min at 37°C. Chromatin was digested using 500U DpnII (NEB) overnight at 37°C while shaking, and the digestion was repeated with an additional 250U of enzymes for 8 hours, meanwhile digestion was determined by gel electrophoresis. Enzyme was deactivated at 65°C for 20 min. Chromatin samples were divided in 3 tubes, diluted and ligated by adding H2O up to 1.2 ml, 133 µl T4 ligase buffer 10X and 6000U total NEB T4 DNA Ligase (M0202M) per tube and incubating at 16°C overnight while shaking. Ligation efficiency was checked by gel electrophoresis. Chromatin was de-crosslinked with proteinase K overnight at 65°C, and treated with RNase A at 37°C for 1 hour. DNA was extracted by

Phenol:Choroform and precipitated with Ethanol. Purified DNA was digested with 50U NEB Csp6I overnight at 37°C while shaking, and digestion was determined by gel electrophoresis. Enzyme was deactivated at 65°C for 20 min. DNA circularization was performed using 4000U NEB T4 DNA Ligase overnight at 16°C. A total of 1 µg DNA was amplified per sample with inverse PCR primers containing Illumina forward and reverse sequencing adapters (see Key resources for sequences). PCR was performed using Expand™ Long Template PCR System (Sigma) with the following thermocycler program: 94°C for 2 min; 94°C for 15 sec; 53-55°C for 1 min; 68°C for 2.30 min; repeat for 29 cycles; 68°C for 7 min; hold at 4°C. 4C-seq libraries were size-selected on gel to remove any potential primer dimers and fragments above 700 bp, then quantified using RT-PCR (KAPA Biosystems) and sequenced using 50bp single-end on Illumina HiSeq 2500.

SYK 4C bait primers:

DpnII restriction site:

AATGATACGGCGACCACCGAGATCTACACTCTTTCCCTACACGACGCTCTTCCGATCTNNNNNNGAGGGCATTCCCATTAGATC (NNNNNN: barcode sequence different for each sample).

Csp6I restriction site:

CAAGCAGAAGACGGCATACGAGATAGGTCGCAGTGACTGGAGTTCAGACGTGTGCTCTTCCGATCTtaatctttggataagtggcc.

**Quantification and Statistical Analysis**

**RNA-Seq Data processing and quality control**

Paired-end reads were mapped to the hg38 genome using TopHat2 [10] (parameters:–no-coverage-search–no-discordant–no-mixed–b2-very-sensitive–N 1). Bigwigs were obtained for visualization on individual as well as merged bam files using Deeptools/2.3.3 [11] (parameters bamCoverage --binSize 1 --normalizeUsing RPKM). Counts for Refseq genes were obtained using htseq-counts [12]. Principal Component Analysis was performed using R to check the reproducibility of replicates (See **Supplementary Figure S1A**). DESeq2 version 1.4 [13] was used to normalize expression counts and get differentially expressed genes (absolute log2 fold-change> 1 and FDR < 0.01). Gene Set Enrichment Analysis of gene expression in NSD2 High versus Low cells were performed using GSEA desktop application on normalized reads counts for each replicate from the 26586 protein coding genes of hg38 genome.

**ATAC-seq Data processing and quality control**

Reads were aligned to hg38 genome with Bowtie2 [14] (parameters: –no-discordant -p 12 –no-mixed -N 1 -X 2000). Potential PCR duplicates were removed from the reads with Picard-tools. ATAC-seq peaks were called with PeaKDEck [15] (parameters: -sig 0.0001 -PVAL ON). Bigwigs were obtained for visualization on individual as well as merged bam files using Deeptools/2.3.3 (parameters bamCoverage --binSize 1 --normalizeUsing RPKM).

**4C-seq Data processing and quality control**

Processing of 4C-seq data was performed using 4Cker pipeline [8]. Briefly, mapping was performed using Bowtie2 to a reduced genome consisting of all unique 24-nt-long regions surrounding DpnII sites from the human reference genome (hg38), allowing for zero mismatches. For comparison between conditions, DESeq2 version 1.4 [13] with default parameters was used to normalize total read count per window between samples and to identify the windows with significant 4C signal differences, using an FDR-adjusted p-value cutoff of 0.05.

**ChIP-seq Data processing and quality control**

Reads were aligned to hg38 genome with Bowtie2 [14] (parameters: –no-discordant -p 12 –no-mixed -N 1 -X 2000). Ambiguous reads were filtered to use uniquely mapped reads in the downstream analysis. PCR duplicates were removed using Picard-tools (version 1.88). Bigwigs were obtained for visualization on individual as well as merged bam files using Deeptools/2.3.3 [11] (parameters bamCoverage --binSize 1 --normalizeUsing RPKM; or bamCompare --verbose --

binSize 25 --ratio subtract --scaleFactorsMethod SES for subtraction files). For H3K36me2 ChIP-seq, an internal Drosophila spike-in was added. Reads were aligned to dm6 genome with Bowtie2 (parameters: –no-discordant -p 12 –no-mixed -N 1 -X 2000). Bigwigs were created after normalization with the spike-in Drosophila read counts. Heatmaps and average profiles were performed on merged bigwig files using Deeptools/2.3.3. For CTCF, H3K27ac and Rad21 ChIP-seq, MACS version 1.4.2 [16] was used to call peaks (parameters: -p 1e-6 -g hs -B --single-profile). For CTCF and H3K27ac ChIP-seq, a reference list of peaks coordinates was created containing all the peaks present in any replicate, and merging overlapping peaks using Bedtools merge –i. Counts for the reference list of peak coordinates were obtained using htseq-counts [12]. PCA was performed using R to check the reproducibility of replicates (See **Supplementary Figure S1B and S1D)**. DESeq2 version 1.4 [13] was used to normalize read counts and get differential peaks (absolute log2 fold-change> 1 and FDR < 0.01).

**Hi-C Data processing and quality control**

**Processing**

HiC-Bench [2] was used to align and filter the Hi-C data, identify TADs, and generate Hi-C heatmaps. To generate Hi-C filtered contact matrices, the Hi-C reads were aligned against the human reference genome (hg38) by bowtie2 [14] (version 2.3.1). Mapped read pairs were filtered by the GenomicTools [17] tools-hic filter command integrated in HiC-bench for known artifacts of the Hi-C protocol. The filtered reads include multi-mapped reads (‘multihit’), read-pairs with only one mappable read (‘single sided’), duplicated read-pairs (‘ds.duplicate’), low mapping quality reads (MAPQ < 30), read-pairs resulting from self-ligated fragments, and short-range interactions resulting from read-pairs aligning within 25kb (‘ds.filtered’). For the downstream analyses, all the accepted intra-chromosomal read-pairs (‘ds.accepted intra’) were used. The Hi-C filtered contact matrices were corrected using the ICE “correction” algorithm [18] built into HiC-bench. TADs and boundaries were identified using the Crane method [19] at 40 kb bin resolution with an insulating window of 500kb. HiC heatmaps for regions of interest were generated using the ICE corrected contact matrices through the 'hic-plotter-diff' pipeline step integrated in HiC-bench.

**Quality Control and TAD / Boundary Stats**

Quality assessment analysis shows that the total numbers of reads in the samples ranged from ~120 million to ~185 million (Supplementary Figure S3A). The percentage of reads aligned was always over 98% in all samples. The proportion of accepted reads ('ds-accepted-intra' and 'ds-accepted-inter') were in the range of ~46 - 47%.

The number of TADs identified by the Crane method across replicates ranged between 3460 and 3532 from which ~500 TADs (14%) showed sizes smaller or equal to 80kb in each replicate. We removed such short length TADs from the following metrics analysis, considering this approach more representative. The distribution of TAD sizes showed that 85% of the TADs ranged between 160 kb and 1.04 Mb with a median of 480 kb in both NSD2 High and Low conditions. The mean TAD sizes were 573 kb and 591 kb for NSD2 High and Low conditions, respectively (Supplementary Figure S3G).

R (prcomp, scale=TRUE and center=TRUE) was used to perform a PCA on the Hi-C datasets using the "ratio" insulation scores produced by HiC-Bench (bins = 40kb) after ICE correction (Supplementary Figure S3B).

**DOWNSTREAM ANALYSIS: TADs, Boundaries, and Compartments**

**Screening of Potentially Altered Boundaries**

The Hi-C downstream analysis involved a genome-wide screening of TAD boundary insulation changes in NSD2 High cells based on boundary insulation scores (ratio index) and CTCF/Rad21 binding enrichment. Individual cases of potentially altered boundaries were confirmed by visual inspection of HiC heatmaps.

**Mean Boundary Insulation Scores (Ratio index)**

To assess and compare boundary strength alteration in NSD2 High versus Low cells, we included the calculation of the Mean Boundary Score (MBS) for every boundary identified in the NSD2 Low condition (reference boundaries). For this purpose, we used the ‘by group’ NSD2 Low TADs identified by the Crane method. HiC-Bench performs a ‘by group’ analysis by merging the DNA interaction information of all the replicates of the same condition, in this case, to identify TADs per condition.

We calculated the MBS as the arithmetic mean of the ‘ratio’ insulation scores inside the reference boundary coordinates being assessed. HiC-Bench calculates one ratio score per bin (40kb) as explained in Lazaris et al. (2017). As a result, there are generally multiple insulation scores per TAD boundary identified by the Crane algorithm. The number of bins inside boundaries showed a median number of 7 and, accordingly, the boundary median size was 280 kb. The MBS was used to calculate NSD2 High MBS logFC values with respect to the NSD2 Low condition. A differential analysis on the ratio insulation scores inside each boundary was also performed. An unpaired t-test (two-sided) was used by pooling all the ratio insulation scores inside the reference boundary coordinates and adjusting with FDR correction.

**CTCF and Rad21 Occupancy in Boundaries: Integration with Insulation Data**

The CTCF and Rad21 peaks were mapped to the boundaries to integrate the boundary insulation data obtained with the CTCF and Rad21 binding data. We assigned a peak to a boundary if the peak overlapped with the boundary region (> 0 bp). An extension of the boundary region by 1 bin (40 kb) on either side of the boundary was considered. The CTCF/Rad21 signal of all the peaks assigned to a boundary were aggregated and then NSD2 High versus Low logFC values were calculated. Significant changes in global CTCF/Rad21 occupancy within the boundaries were calculated using a two-sided unpaired t-test by pooling all the CTCF/Rad21 peak intensities assigned in the boundary.

A ranked-table of boundary coordinates with the insulation and CTCF/Rad21 metrics was created. To generate Hi-C heatmaps, the best-ranked boundary cases were selected by taking into account unidirectional significant fold changes in MBS, CTCF and Rad21 (FDR < 0.01). In this approach, we assumed that there would be a positive correlation between boundary insulation scores and CTCF/Rad21 binding in boundaries. In addition, we screened each boundary candidate and the adjacent TADs and boundaries in search of significant deregulated genes, gain / loss of interactions and significant changes in H3K27ac.

**Compartments**

Compartment analysis was carried out using the Homer pipeline (v4.6) [19]. Homer performs a principal component analysis of the normalized interaction matrices and uses the PC1 component to define regions of active (A compartments) and inactive chromatin (B compartments). HiC filtered matrices were given as input to run Homer with default parameters (50kb resolution). To confirm the proper sign of the A and B compartment, we used the --active parameter to input peaks of the active mark H3K27ac. We use Homer to compare the interaction profiles in both experiments and calculate a correlation. If one region interacts similarly with other regions in both conditions then the correlation will be high. On the other hand, the correlation will be low if a locus interacts with different regions in both conditions. Using Homer's getHiCcorrDiff.pl, we compared the interaction profiles of both conditions and obtained a correlation difference to identify stable and switching compartments. Altered compartments are named as AB and BA for regions switching from A to B and B to A, respectively.

**Intra-TAD interactions**

To assess statistically significant intra-TAD interactions, we used an algorithm developed by Andreas Kloetgen. As a first step, the algorithm identifies overlapped or positionally consistent TADs (common TADs). This approach establishes a minimum TAD length of 10 bins (400kb) and extends either side of the TAD by 3 bins (+/-120 kb in 40kb resolution). TADs across two samples are considered positionally consistent if their boundaries are as close as 3 bins. The boundaries of the common TAD are then set to those that yield the largest TAD. The set of common TADs between any two samples $s_1$ and $s_2$ is denoted as T. In the next step, a paired two-sided t-test is performed on each single interaction bin within each common TAD between the two samples. It calculates the difference between the average scores of all interaction intensities within such TADs. A multiple testing correction by calculating the false-discovery rate per common TAD (using the R function p.adjust with method="fdr") is also calculated.

$$(1)\ TAD\ interactions\ change(t) = \left(\frac{\sum_{i\in I_t} s_{2_i}}{\# I_t}\right) - \left(\frac{\sum_{i\in I_t} s_{1_i}}{\# I_t}\right)$$

for each $t \in T$, and $I_t$ being all intra-TAD interactions for TAD t.

We classified the common TADs in terms of Loss, Gain or Stable intra-TAD interactions by using FDR < 0.1 and absolute TAD interactions change > 0.3.

**Data Integration: Global Analysis.**

**AB and BA Compartment Changes.**

To show the correlation between the different measurements and the compartment changes**,** we mapped the peaks obtained in H3K27ac2, CTCF, RAD21 and ATAC-seq to the AB, BA and Stable regions. We used the peak intensity values to calculate the peak intensity fold change between NSD2 High and Low and the mean fold change of all the peaks assigned to a compartment region.

Similarly, in the case of RNA-seq data, genes were mapped to the compartment regions and the mean fold change of all the genes assigned to a region was calculated by using the DESeq2 fold change data.

We assigned a peak or gene to a compartment region when the complete peak or gene coordinate was found inside the compartment coordinates. For sparse measurements as in H3K36me2 and H3K27me3 chromatin marks, we used HTSeq to obtain read counts on the AB, BA and Stable regions. Next, we used DESeq2 to normalize the read counts across NSD2 High and Low replicates and to obtain fold change values.

To show the correlation between the different measurements and the compartment changes, we used the mean fold change values to generate boxplots. Statistical significance was assessed using a paired two-sided Wilcoxon rank-sum test.

**Intra-TAD Interactions. Gain, Loss and Stable TADs.**

We used the same method as described in the previous subsection to integrate the H3K27ac2, CTCF, RAD21, ATAC-seq, RNA-seq data, H3K36me2, H3K27me3 measurements with the different intra-TAD interactions subgroups described above (Gain, Loss and Stable TADs).

**Data Integration: Local Analysis in Common TADs.**

**CTCF binding, H3K27ac and RNA expression.**

To compute the total number of differential changes in CTCF binding, RNA expression, and H3K27ac mark that fall within common TADs. We overlapped all CTCF peaks, genes, and H3K27ac peaks with common TADs. Volcano plots were generated using the log2 fold changes and –log10 (p-value) for all three data types (CTCF-ChIP, RNA-seq, H3K27ac-ChIP). Overlapping features with log2 fold change greater than 1 and a q-value less than 0.01 were colored red, while overlapping features with log2 fold change less -1 and a q-value less than 0.01 were colored in blue, and all others were colored in black.

To assess the correlation of CTCF binding, H3K27ac, and RNA expression inside each Common TAD, we assigned a peak or gene to each Common TAD when the complete peak or gene coordinate was found inside the TAD coordinates. Common TADs lacking either CTCF, H3K27ac, or RNA expression features were filtered.

The mean fold change of each feature (CTCF, H3K27ac and RNA expression) inside each Common TAD was computed by two methods. One method considered only the differential peaks / genes found inside a Common TAD (FDR < 0.05) in the mean fold change calculation. The second method considered all the peaks / genes assigned to a Common TAD.

**Intra-TAD Interactions and Compartment Alteration.**

One mean log fold change value for each feature (CTCF, H3K27ac and RNA expression) was assigned to every Common TAD, together with the intra-TAD interactions fold change value previously calculated (see '**Intra-TAD interactions'** subsection). Compartment alteration was also assessed by calculating the mean PC1 value of each Common TAD (50kb bins) and computing the PC1 mean difference between conditions (NSD2 High – NSD2 low). A positive value of the PC1 mean difference indicates that in the NSD2 High condition, that Common TAD had become more active. The higher the PC1 mean difference the stronger the compartment alteration change. To confirm if the PC1 mean difference correlated with real compartment changes, we looked at whether the significant AB and BA regions identified by Homer overlapped with the Common TADs. We then computed the overlap length.

To show the intra-TAD association of all the five features (CTCF, H3K27ac, RNA expression, Intra-TAD interactions and PC1 mean difference) we classified the Common TADs in 'positive', 'negative' or 'no correlated' groups, by taking into account the direction of all the five features (positive correlation). The Pearson Correlation Coefficient (PCC) was calculated by using R ('Cor' and 'pairs' command). 3D plots were also generated using R ('plot_ly' command of the 'plotly' library).

**HiChIP Data processing and quality control**

HiChIP paired-end reads were aligned to hg38 genome using the HiC-Pro pipeline [20]. Default settings were used to remove duplicate reads, assign reads to MboI restriction fragments, filter for valid pairs, and generate 10kb binned interaction matrices. FitHiChIP [21] was used to identify statistically significant chromosomal interactions from the CTCF HiChIP experiments, using a bin size of 10KB. FitHiChIP allows users to either infer peaks from their HiChIP data or use a reference list of peaks from ChIP-seq data. Statistically significant HiChIP interactions ($q<0.01$) were called separately in each experiment with a minimum distance of 20kB and a maximum distance of 3MB. The UseP2PBackgrnd parameter was set to 1 for peak to peak background estimation. HiChIP interactions were considered statistically significant if a peak was found in at least one anchor (Peak to all interactions). This resulted in 8,651 statistically significant CTCF peak to all interactions in NSD2 High and 4,914 CTCF peak to all interactions in NSD2 low. Common CTCF high confidence loops in NSD2 high and NSD2 low were identified if both pairs of loop anchors are overlapping. The median size for common loops was calculated to be 210kB. Boxplots were generated to show the mean contact counts for common CTCF loops in NSD2 low and NSD2 high experiments.

**HICHIP Data integration with CTCF ChIP-seq, RNA-seq and H3K27ac ChIP-seq**

To compute the total number of differential changes in CTCF binding, RNA expression, and H3K27ac marks that fall within common CTCF HiChIP loops, we overlapped all CTCF peaks, genes (5kb upstream of TSS), and H3K27ac peaks within common HiChIP loops (full loop plus anchors). Volcano plots were generated using the log2 fold changes and –log10 (p-value) for all three data types (CTCF-ChIP, RNA-seq, H3K27ac-ChIP). Overlapping features with log2 fold change greater than 1 and a q-value less than 0.01 were colored red, while overlapping features with log2 fold change less -1 and a q-value less than 0.01 were colored in blue, and all others were colored in black.

To assess the correlation of CTCF binding, H3K27ac, and RNA expression inside each Common CTCF Loops, we applied the same approach used to assess the correlation of these features in Common TADs. First, all CTCF peaks were overlapped with all the common loops, and the mean log2 fold change for all peaks found within a common loop were calculated. The same approach was applied to calculate the mean log2 fold change for all genes and for all H3K27ac peaks within each common loop.

To correlate the mean log2 fold changes of all three features, we considered only the common loops that had at least one overlapping differential CTCF and H3K27ac peak as well as one gene expression change. Pairwise correlation plots were generated for the three comparisons, so that each individual point in the scatterplot represents a common CTCF loop. Loops that had positive mean log2 fold change for all three features were colored in red, while loops that had negative log2 fold change for all three features were colored in blue.

To determine if there is a correlation between the significant alterations of CTCF, RNA expression, and H3K27ac, we performed the same analysis, but first filtered for significant changes in CTCF, RNA expression, and H3K27ac peaks, with a q-value of less than 0.05 and an absolute value of the log2 fold change greater than 1. Next, the mean log 2 fold change was computed for all the differential CTCF, gene expression changes, and H3K27ac peaks that overlapped common HiChIP loops. Pearson correlation was calculated for all three comparisons.

**Logistic regression model**

To evaluate which factors contribute most to gene regulation, we modeled the probability that a gene is differentially expressed as a function of chromatin features in insulated domains (TADs and/or CTCF loops) using logistic regression. The covariates of our model were four binary variables which indicate whether each gene shares a CTCF HiChIP loop or a TAD with at least one differential CTCF or H3K27ac peak (data.frame = Supplementary Table 4). The model was fitted using R's glm function as following:

glm(gene.de ~ k27ac.tad + ctcf.tad + k27ac.loop + ctcf.loop, family = binomial(), data = data.frame). The standard Wald-test was used to access the significance of each factor.

### Supplementary Figure 1

**a** Principal Component Analysis of RNAseq replicates

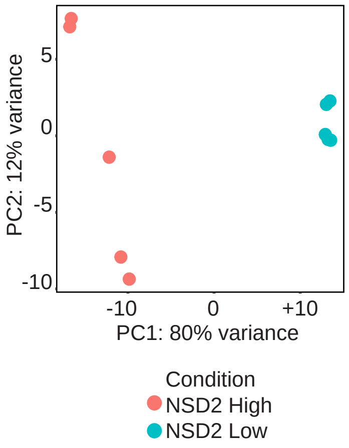


**b** Gene set enrichment analysis of RNAseq in NSD2 High versus Low cells

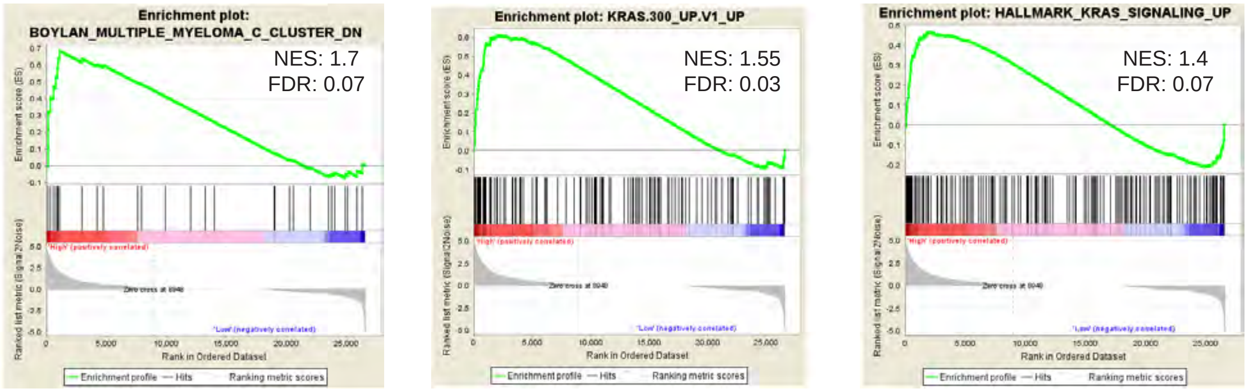

### Supplementary Figure 2

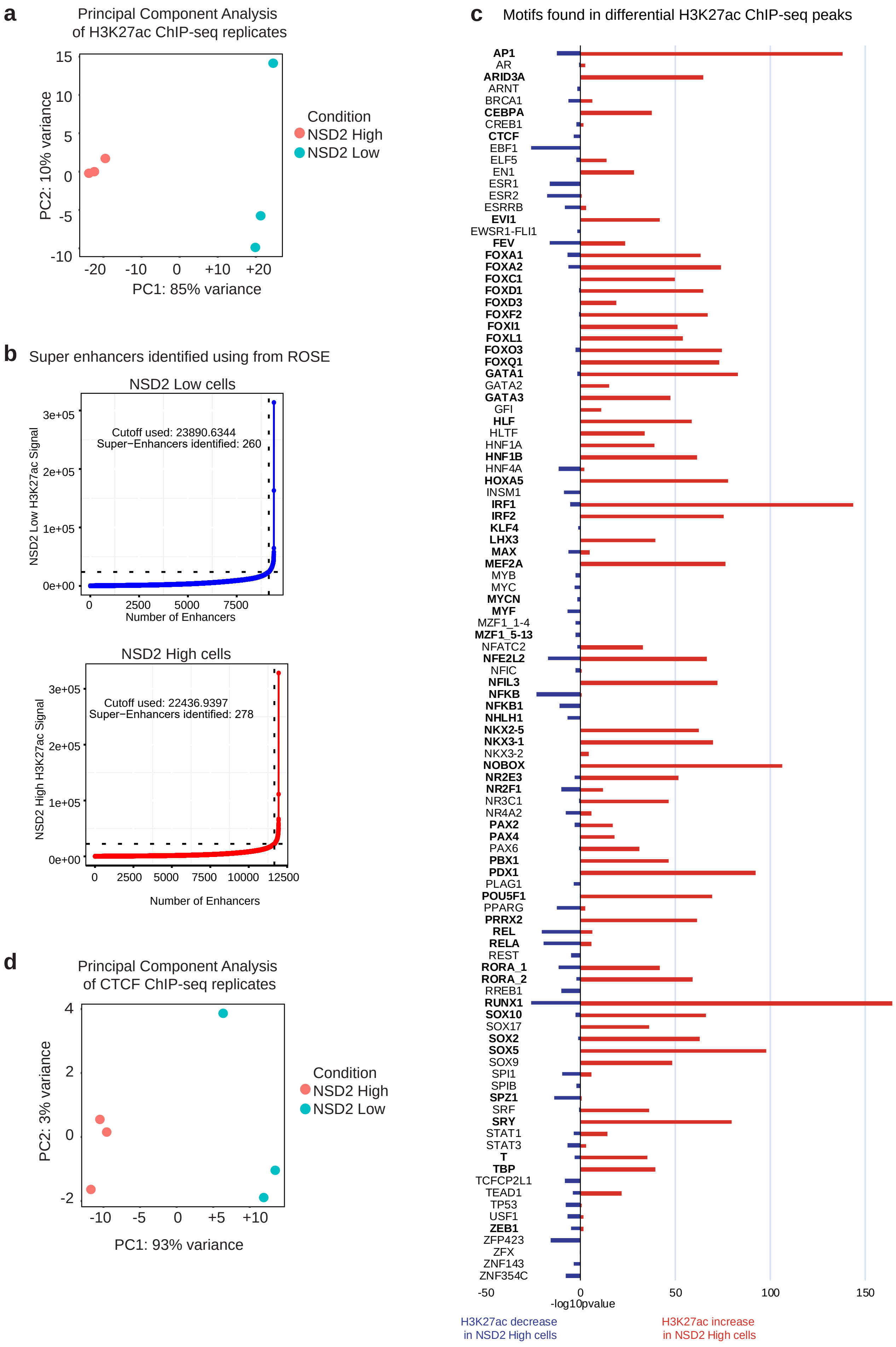

### Supplementary Figure 3

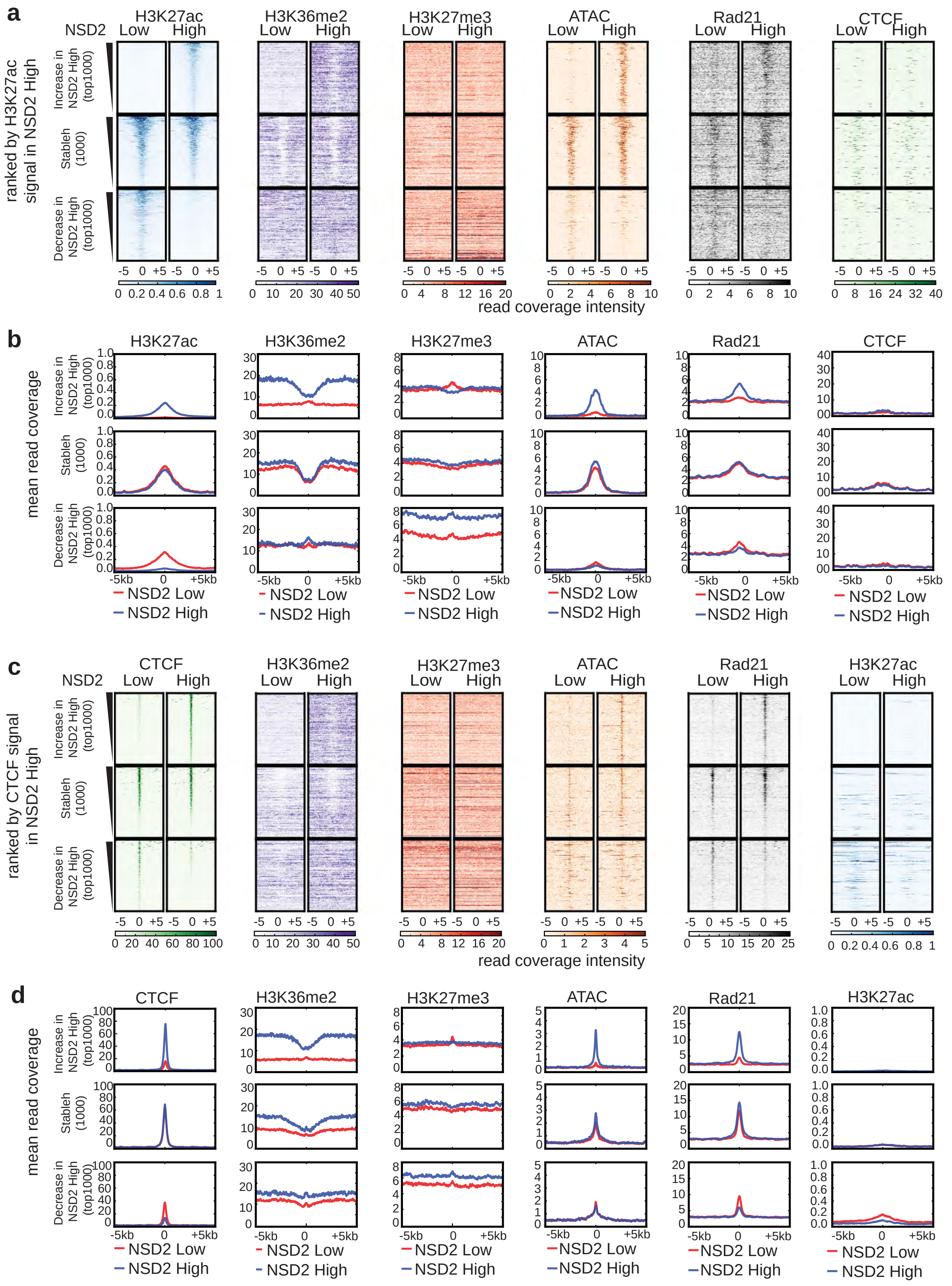

### Supplementary Figure 4

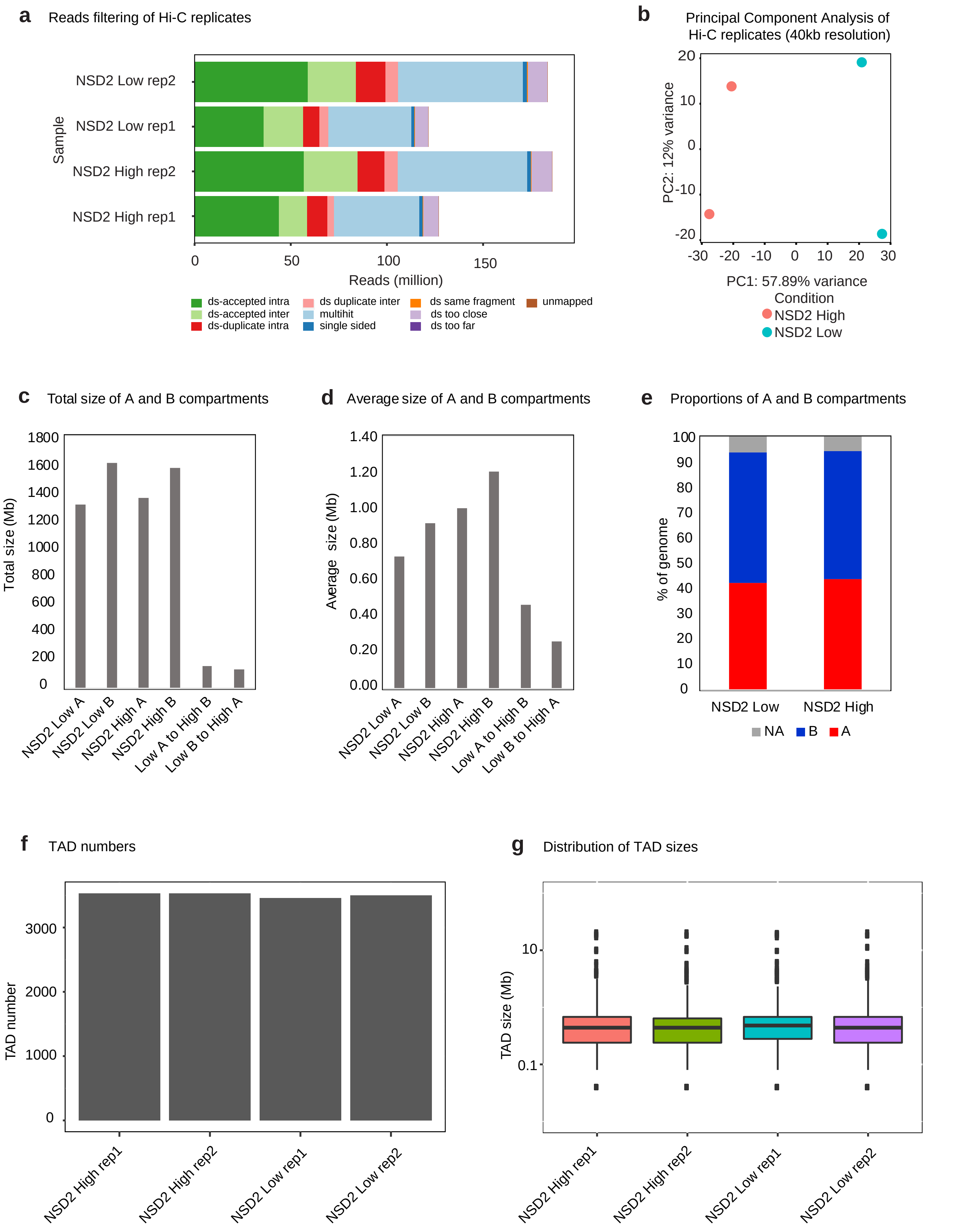

### Supplementary Figure 5

**a** Example of altered Hi-C surrounding the *FGF13* gene

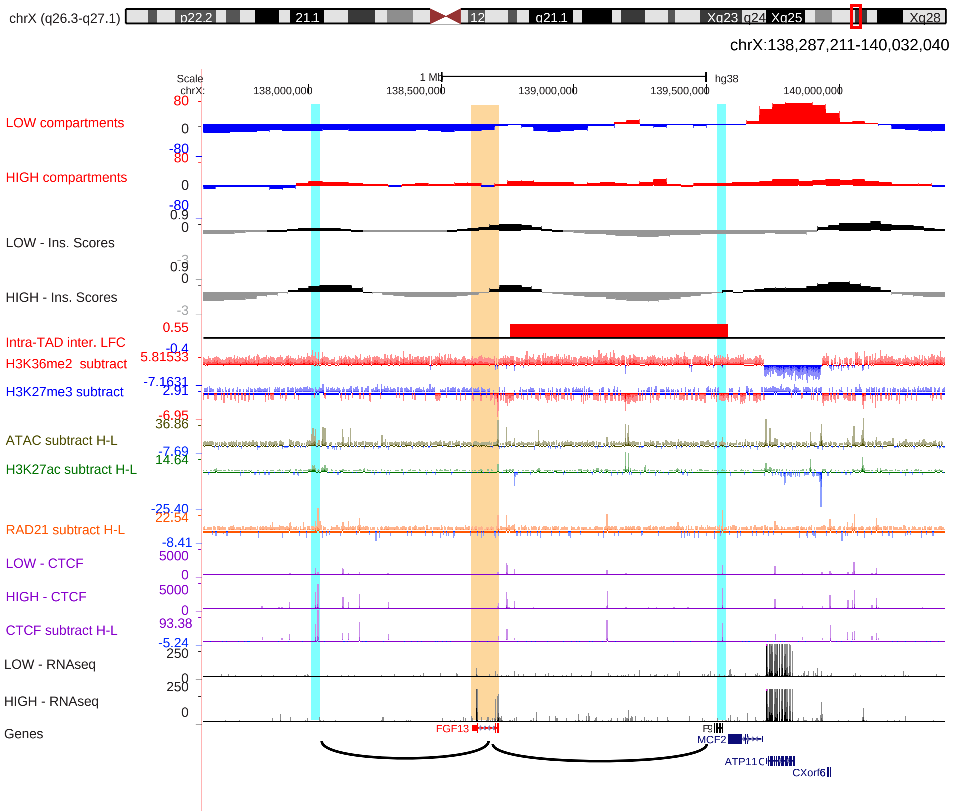


**b** Hi-C plots of the region surrounding the *FGF13* gene

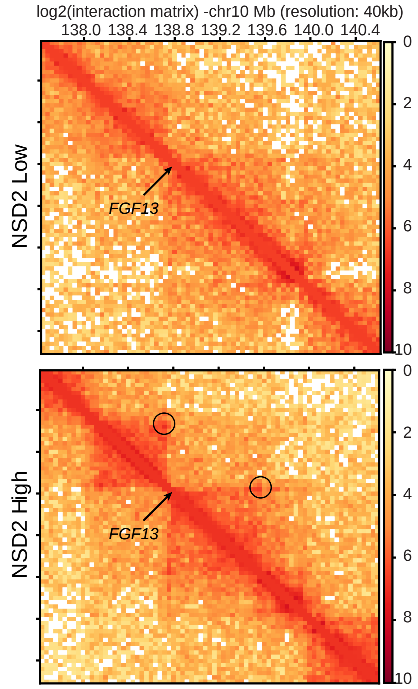


**c** UCSC tracks showing chromatin features in the region surrounding the *KRAS* gene

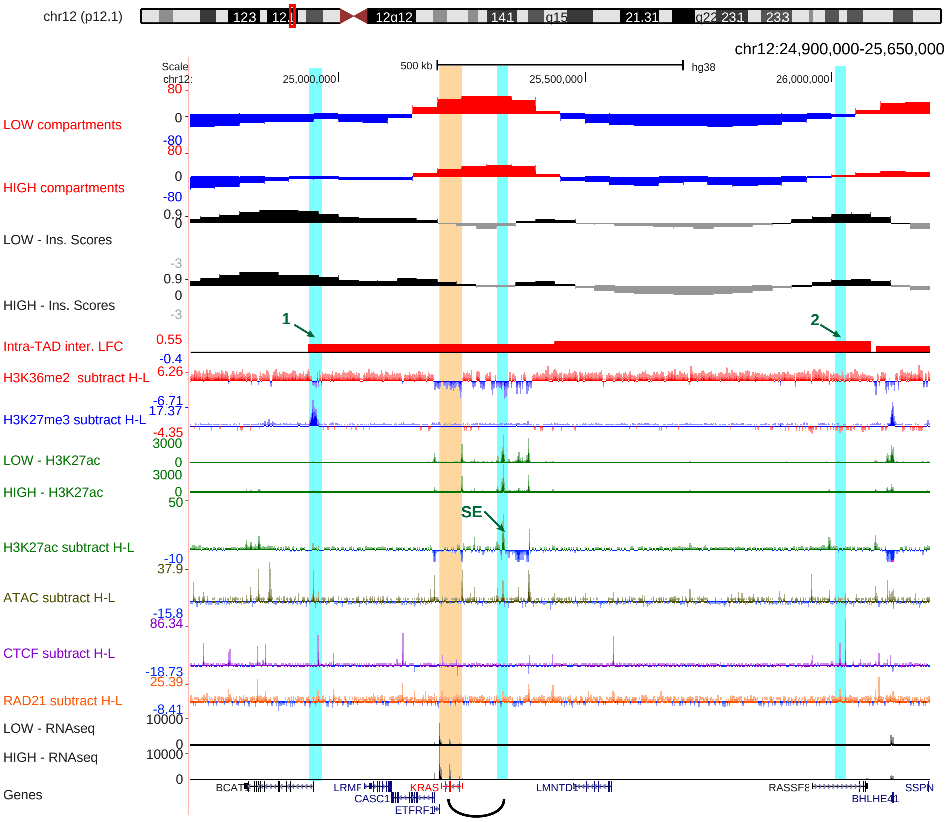


**d** Hi-C plots of the region surrounding the *KRAS* gene

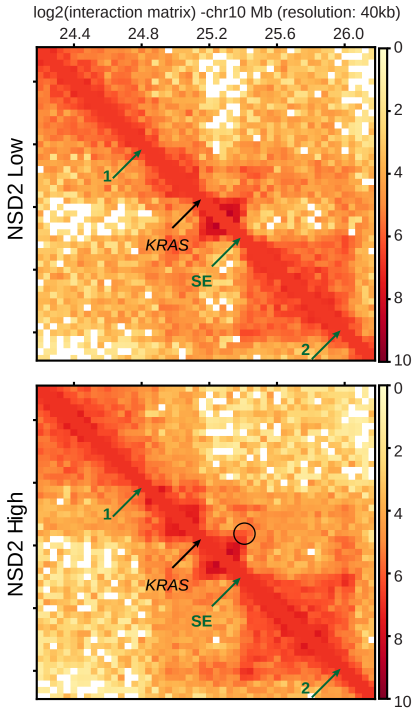

### Supplementary Figure 6

**a** Significant changes in gene expression, CTCF and H3K27ac peaks in TADs and CTCF HiChIP loops

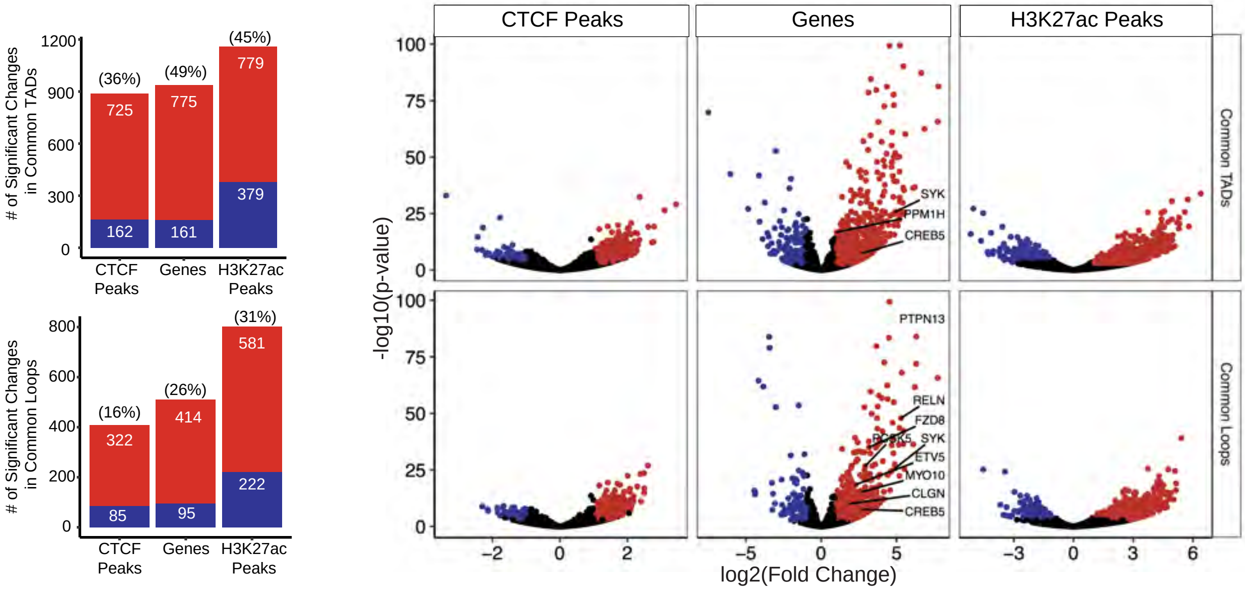


**b**

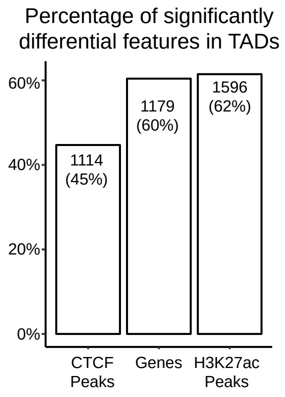


**c**

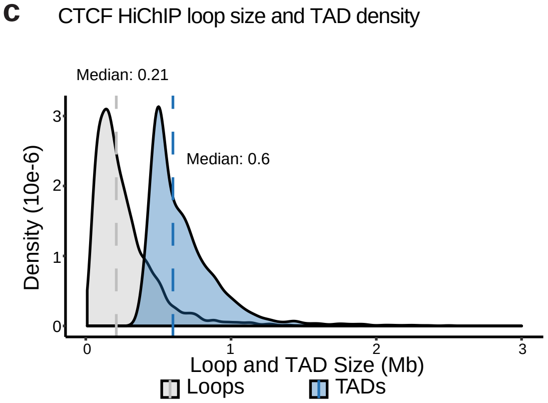

### Supplementary Figure 7

**a** Correlation of gene expression, H3K27Ac, CTCF intra-TAD interactions and compartments within TADs

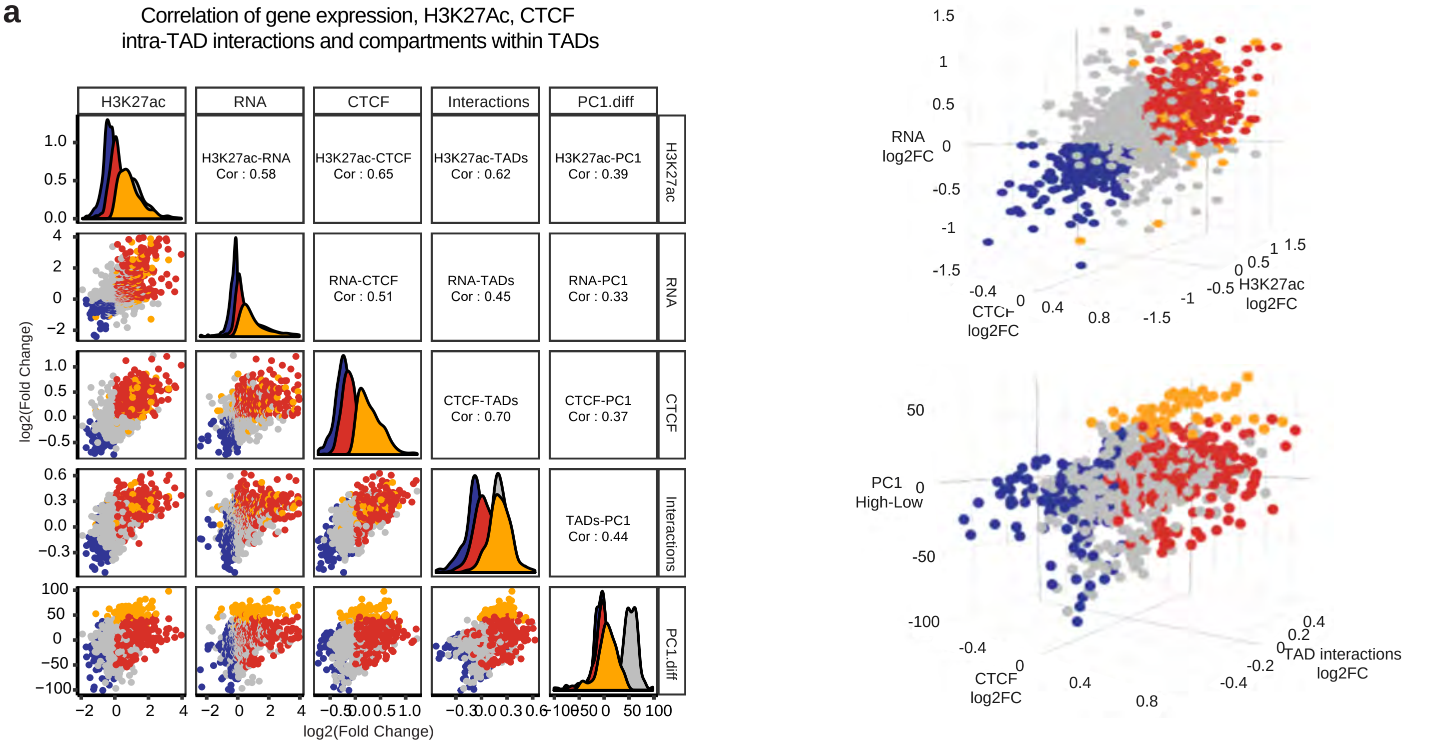


**b** Correlation of expression, H3K27ac and CTCF changes within CTCF HiChIP loops

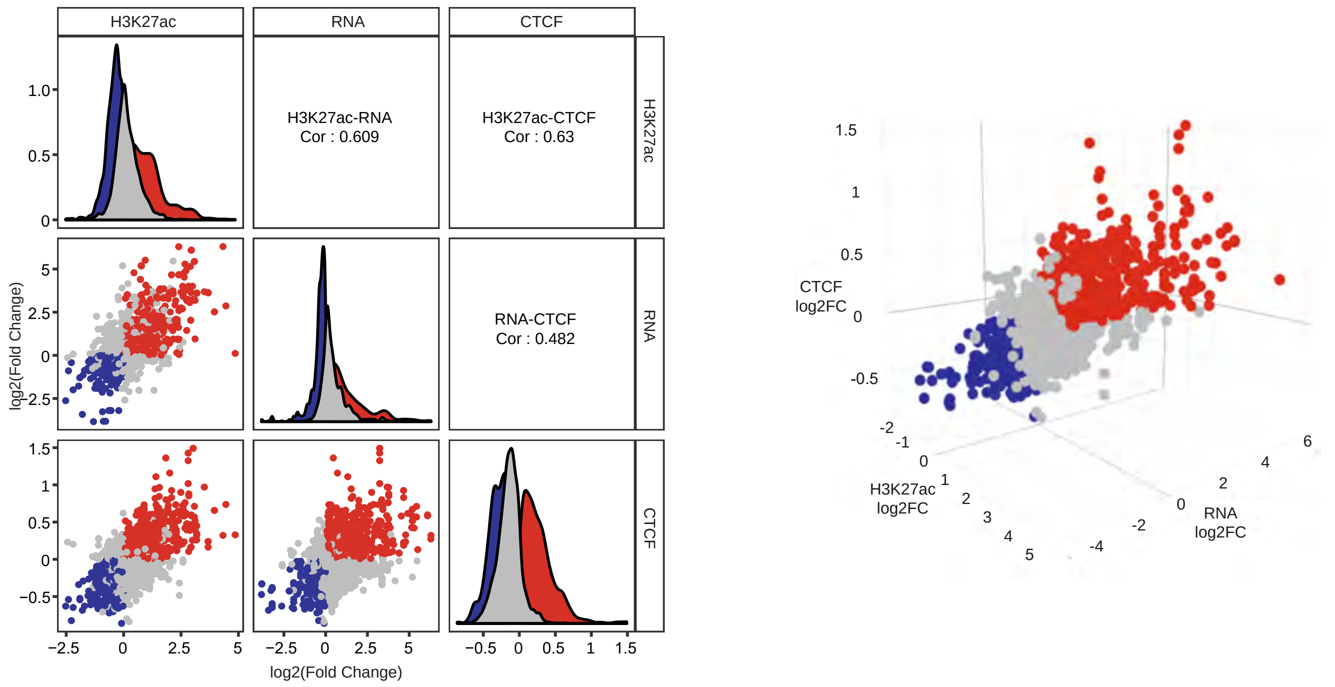


**c** Concordant and disconcordant changes in gene expression, H3K27Ac and CTCF within TADs and CTCF HiChIP loops

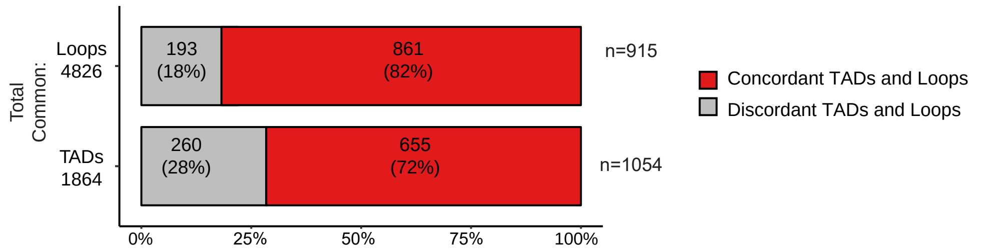
